# Mechanism of pharmacochaperoning in KATP channels revealed by cryo-EM

**DOI:** 10.1101/572297

**Authors:** Gregory M. Martin, Min Woo Sung, Zhongying Yang, Laura M. Innes, Balamurugan Kandasamy, Larry L. David, Craig Yoshioka, Show-Ling Shyng

## Abstract

ATP-sensitive potassium (K_ATP_) channels composed of a pore-forming Kir6.2 potassium channel and a regulatory ABC transporter sulfonylurea receptor 1 (SUR1) regulate insulin secretion in pancreatic β-cells to maintain glucose homeostasis. Mutations that impair channel folding or assembly prevent cell surface expression and cause congenital hyperinsulinism. Structurally diverse K_ATP_ inhibitors have been shown to act as pharmacochaperones to correct mutant channel expression, but the mechanism is unknown. Here, we compare cryoEM structures of K_ATP_ channels bound to pharmacochaperones glibenclamide, repaglinide, and carbamazepine. We found all three drugs bind within a common pocket in SUR1. Further, we found the N-terminus of Kir6.2 inserted within the central cavity of the SUR1 ABC core, adjacent the drug binding pocket. The findings reveal a common mechanism by which diverse compounds stabilize the Kir6.2 N-terminus within the SUR1 ABC core, allowing it to act as a firm “handle” for the assembly of metastable mutant SUR1-Kir6.2 complexes.

## Introduction

ATP-binding cassette transporters (ABC transporters) comprise a large protein superfamily responsible for transporting diverse molecules across cell membranes (Thomas and Tampe, 2018; Trowitzsch and Tampe, 2018; Wilkens, 2015). Uniquely, the sulfonylurea receptors SUR1 and SUR2 lack transport activity *per se* but instead are devoted to forming the ATP-sensitive potassium (K_ATP_) channels, wherein four SUR.x subunits form a complex with four subunits of an inwardly rectifying potassium channel, Kir6.1 or Kir6.2 (Aittoniemi et al., 2009; Ashcroft and Ashcroft, 1990; Bryan et al., 2007; Nichols, 2006). K_ATP_ channels are gated by intracellular ATP and ADP, key features which enable them to regulate membrane excitability according to the energetic state of the cell (Nichols, 2006). Expressed in many cell types, they have critical roles in endocrine, cardiovascular, neurological and muscular functions (Aguilar-Bryan and Bryan, 1999; Foster and Coetzee, 2016; Nichols et al., 1996). Critically in pancreatic β-cells, K_ATP_ channels composed of SUR1 and Kir6.2 couple glucose metabolism to insulin secretion (Aguilar-Bryan et al., 2001; Ashcroft, 2005). Gain-of-function mutations in the SUR1 gene *ABCC8* or the Kir6.2 gene *KCNJ11* are a major cause of neonatal diabetes, while loss-of-function mutations result in congenital hyperinsulinism (Ashcroft et al., 2017; Koster et al., 2005; Stanley, 2016). Among the latter, severe hyperinsulinism requiring pancreatectomy to prevent life-threatening hypoglycemia is often the result of K_ATP_ channel mutations that reduce K_ATP_ channel density on the β-cell surface by impairing channel biogenesis, assembly, and trafficking (referred to as trafficking mutations hereinafter) (Huopio et al., 2002; Vajravelu and De Leon, 2018).

Previously, we showed that K_ATP_ channel inhibitors commonly used to treat Type 2 diabetes are often efficient pharmacochaperones for trafficking-impaired mutant K_ATP_ channels, including sulfonylureas (SUs) and glinides (Yan et al., 2004; Yan et al., 2007). Most trafficking mutations identified to date are in SUR1, perhaps due to its large size compared to Kir6.2 (Martin et al., 2013; Snider et al., 2013). The SUR1 protein has an N-terminal transmembrane domain referred to as TMD0 that interacts directly with the Kir6.2 subunit (Babenko and Bryan, 2003; Chan et al., 2003; Lee et al., 2017; Li et al., 2017; Martin et al., 2017b; Schwappach et al., 2000), followed by a cytoplasmic loop termed L0, and then the ABC core structure comprising two transmembrane domains TMD1 and TMD2 and two nucleotide binding domains NBD1 and NBD2 (Aguilar-Bryan et al., 1995; Martin et al., 2017b). Of particular interest, while trafficking mutations are found throughout SUR1, SUs and glinides have to date only corrected defects caused by mutations located in TMD0, the domain contacting Kir6.2. Recently, we found that carbamazepine (CBZ), an anticonvulsant structurally not related to SUs and glinides, also corrects K_ATP_ channel trafficking defects (Chen et al., 2013b; Sampson et al., 2013) and surprisingly is effective only for mutations in TMD0 of SUR1 (Devaraneni et al., 2015; Martin et al., 2016). Furthermore, like SUs and glinides, CBZ inhibits K_ATP_ channel activity by reducing channel *P*_*o*_ and abolishing channel responsiveness to MgADP (Zhou et al., 2014). The striking similarities of SUs, glinides and CBZ in their effects on the channel despite their chemical uniqueness suggest a shared pharmacochaperoning mechanism which remains elusive.

In this study, we took a comparative structural approach to understand how glibenclamide (GBC, a sulfonylurea), repaglinide (RPG, a glinide), and CBZ, interact with the channel to affect channel biogenesis and function. Using single-particle cryo-electron microscopy (cryoEM), we determined structures of the channel in the presence of GBC, RPG, and CBZ, and without pharmacochaperone. The structures show that like GBC (Martin et al., 2017a; Wu et al., 2018), RPG and CBZ occupy the same binding pocket in the transmembrane bundle above NBD1 of SUR1. Further, we undertook structural, biochemical and functional studies to determine the involvement of the distal N-terminal 30 amino acid stretch of Kir6.2, which is functionally critical to the actions of these drugs but whose structure has previously remained unresolved. The evidence from these combined efforts establish that the distal Kir6.2 N-terminus is located in the cavity formed by the two halves of the SUR1 ABC core and is adjacent to the drug binding pocket. The results reveal how a chemically diverse set of K_ATP_ channel inhibitors allosterically control channel gating, and also promote the assembly and trafficking of nascent channels to the β-cell surface, by stabilizing a key labile regulatory interaction between the N-terminus of Kir6.2 and the central cavity the ABC core of SUR1.

## Results

### Structure determination

For structure determination, a FLAG-tagged hamster SUR1 and a rat Kir6.2 (95% and 96% identical to human, respectively) were overexpressed in the insulinoma cell line INS-1, and the channel complex affinity purified via the FLAG-epitope tag as described for our recent cryoEM structure determination of K_ATP_ bound to GBC and ATP (Martin et al., 2017a; Martin et al., 2017b). These channels have been used extensively for structure-function and pharmacochaperone studies and are thus well characterized (Inagaki et al., 1995; Shyng et al., 1998; Yan et al., 2004). To allow direct comparison of channel structures containing bound RPG or CBZ with the GBC/ATP-bound structure solved previously (Martin et al., 2017a), data were similarly collected in the presence of ATP (1mM without Mg^2+^) but varying the pharmacochaperone by alternatively including 30μM RPG (referred to as the RPG/ATP state), 10µM CBZ (referred to as the CBZ/ATP state) or the drug vehicle 0.1% DMSO (referred to as the ATP-only state) in the sample. Further, as an additional control we determined the structure of channels without any pharmacochaperone or ATP (referred to as the apo state).

Initial data processing in RELION with C4 symmetry imposed yielded one dominant 3D class for each dataset, with an overall reported resolution ranging from ∼4Å for the RPG/ATP and CBZ/ATP states to ∼7Å for the ATP-only and the apo states (Fig.1; Fig.1-figure supplements 1-4; Table 1). As with the GBC/ATP state structures we reported previously (Martin et al., 2017a; Martin et al., 2017b), increased disorder was observed at the periphery of the complex in all structures possibly due to minor deviations from C4 symmetry. To see whether resolution of SUR1 could be improved, focused refinement using symmetry expansion and signal subtraction with different masks was performed on all structures, including the previously collected GBC/ATP dataset (Martin et al., 2017a), as described in Methods. This yielded maps with improved reported resolutions for SUR1 as follows: 3.74 Å for the GBC/ATP state, 3.65 Å for the RPG/ATP state, 4.34 Å for the CBZ/ATP state, 4.50 Å for the ATP-only state, and 4.55 Å for the apo state using the FSC cutoff of 0.143 (Table 1).

**Table 1.**
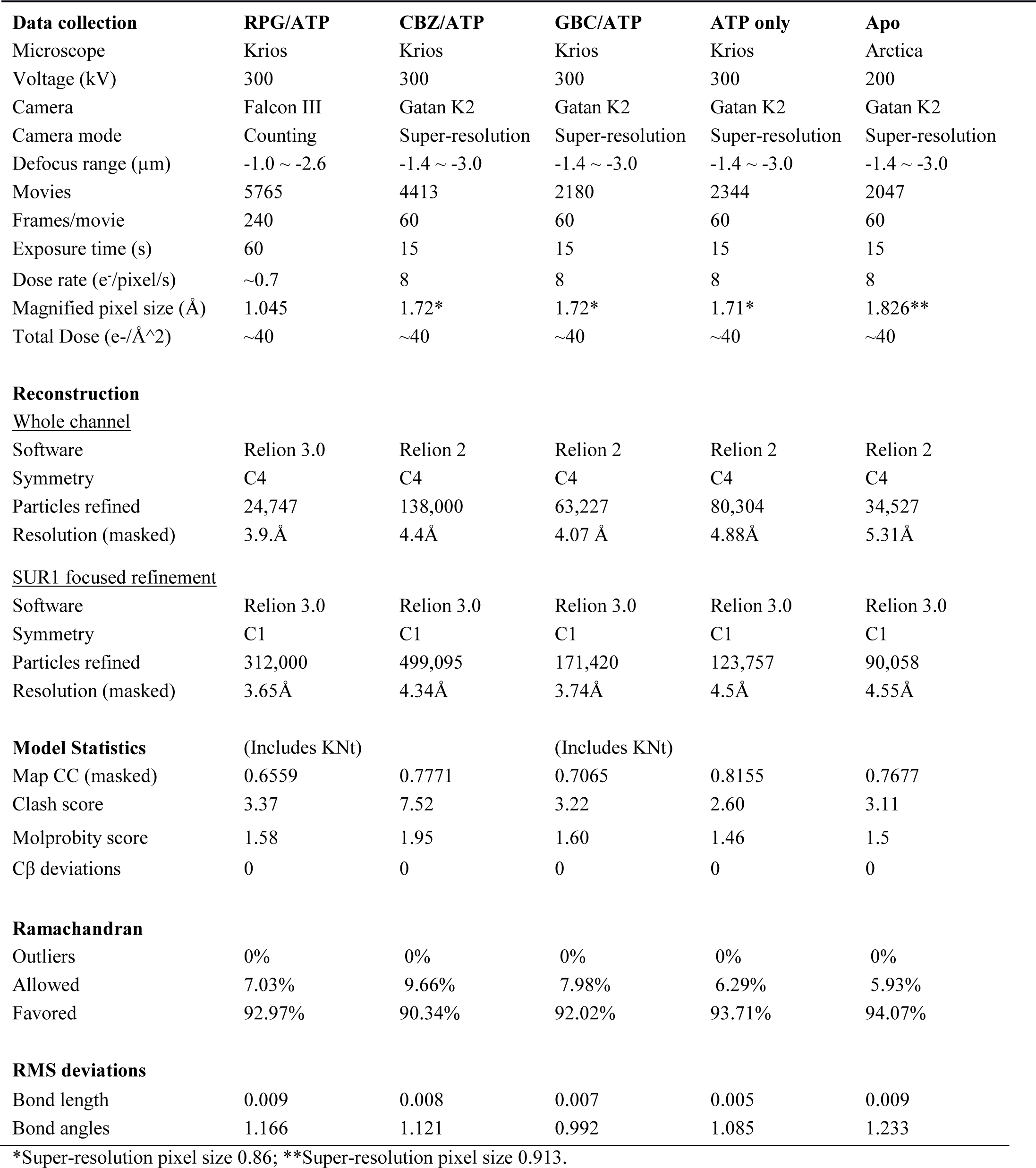
Statistics of cryo-EM data collection, 3D reconstruction and model building.

**Figure 1:**
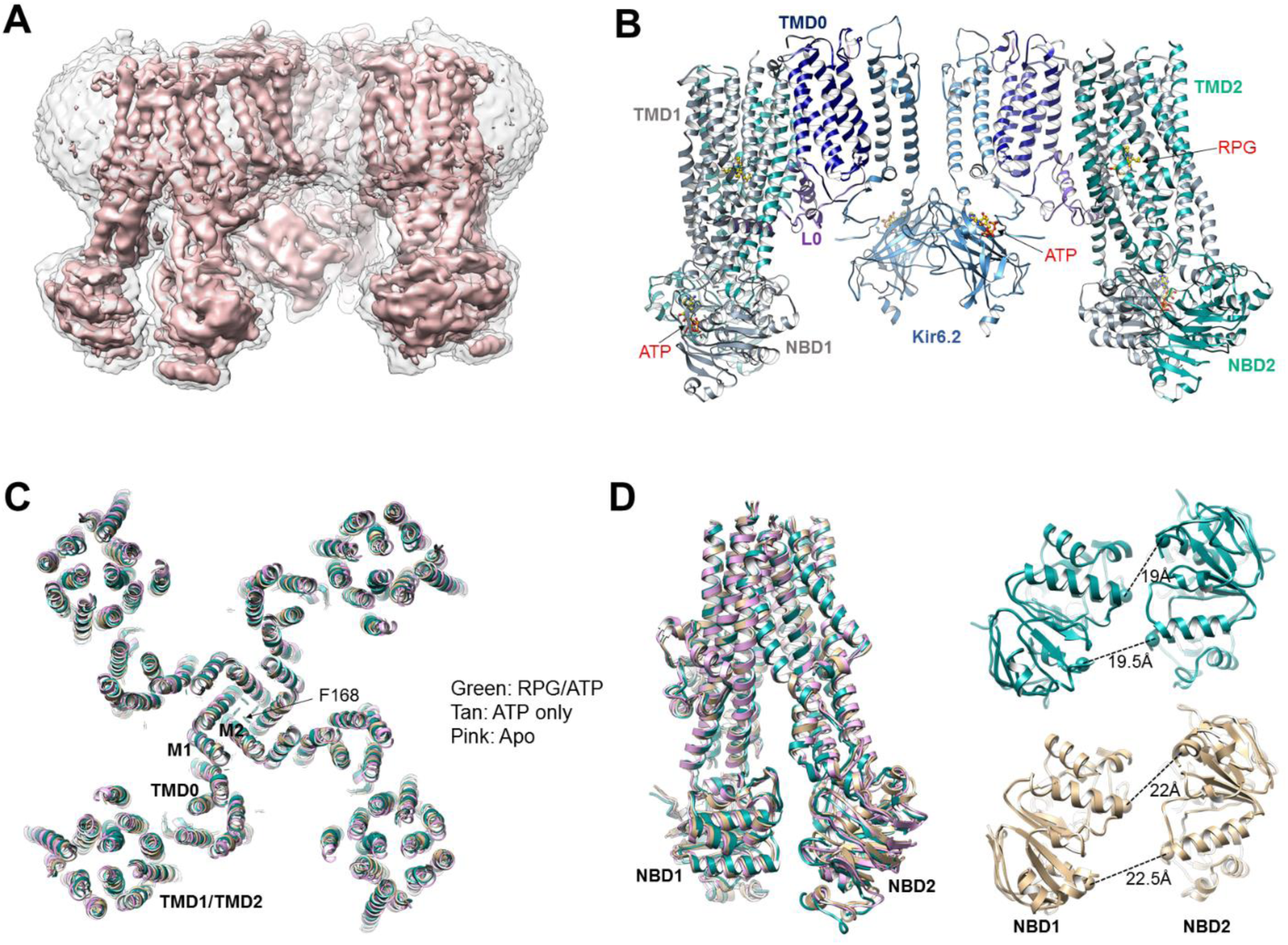
Structural determination and comparison. **(A)** Unsharpened 3.9Å C4 cryoEM reconstruction of the K_ATP_ channel bound to RPG and ATP. **(B)** Structural model of the channel in the RPG/ATP state. **(C)** Overlay of the RPG/ATP state structure, the ATP only state structure, and the apo state structure viewed from the top showing similarity of the dominant class of the ATP only and the apo state to the RPG/ATP state structure. **(D)** *Left*: Same model as **(C)** viewed from the side and focusing on the ABC transporter core module of SUR1 to illustrate the inward-facing conformation observed in all three structures. *Right*: Separation between Walker A and the signature motif in NBD1 and NBD2 (G716::S1483 and S831::G1382; Ca to Ca, indicated by the dashed line) in SUR1 bound to RPG and ATP (green) and ATP only (tan) viewed from the bottom.

For model building, we used our previously published GBC/ATP structure (PDB: 6BAA) (Martin et al., 2017a) as a starting point and refined the models against the experimental data. For the GBC/ATP structure, the new map derived from focused refinement of the Kir6.2 tetramer plus an SUR1 subunit showed significantly improved cryoEM density in a number of regions of SUR1 previously not modeled in the GBC/ATP map (see PDB and EMD files to be deposited). This allowed us to build additional residues into the GBC/ATP structure. In particular, the ATP density in NBD1 became clearly resolved (Fig.1-figure supplement 5A; note ATP binds NBD1 in the absence of Mg^2+^ (Ueda et al., 1997)). The linker between TMD2 and NBD2 (L1319-Q1342) also showed much improved density, allowing us to build a continuous polyalanine model (Fig.1-figure supplement 5A). Models for the other structural states were similarly built according to the highest resolution maps available for the focus-refined regions (for details see Materials and Methods).

In all structures determined in the presence of pharmacochaperones and ATP, the Kir6.2 tetramer is in a closed conformation (Fig.1C), and the SUR1’s ABC core in an “inward-facing” conformation wherein the NBD1 and NBD2 are separated (Fig.1D). This overall structure is very similar to that previously reported for the GBC/ATP state (Martin et al., 2017a; Martin et al., 2017b). Interestingly, the dominant class emerging from 3D classification for the ATP-only state as well as the apo state presented a similar conformation, with Kir6.2 tetramer closed and SUR1 inward-facing (Fig.1C and D). Of note, a dominant inward-facing conformation in the absence of ligand was also observed in another ABC transporter, the multidrug resistance protein MRP1 (Johnson and Chen, 2017). These observations suggest the closed channel conformation is the most stable for the apo state under our experimental conditions (i.e. no ATP and no exogenous PIP_2_, a lipid required for K_ATP_ channel opening (Nichols, 2006)). More data and extensive analyses will be needed to resolve whether other minor conformations are present in our samples and to understand the dynamics of these structures, especially in the ATP-only and the apo states. Here, we focus on analyzing the pharmacochaperone binding pocket and the mechanisms by which pharmacochaperones affect channel assembly and gating.

### RPG and CBZ are located in the GBC binding pocket

We previously solved the K_ATP_ structure in the presence of GBC and ATP, revealing that GBC is lodged in the TM bundle above NBD1 (Martin et al., 2017a), a finding which has been independently confirmed (Wu et al., 2018). Here, in both the RPG/ATP and CBZ/ATP structures, we observe strong and distinctly shaped cryo-EM densities within the same GBC binding pocket (Fig. 2B-D). Such density is absent in the ATP-only and the apo structures (Fig. 2E and 2F), supporting assignment as the pharmacochaperone ligand.

**Figure 2:**
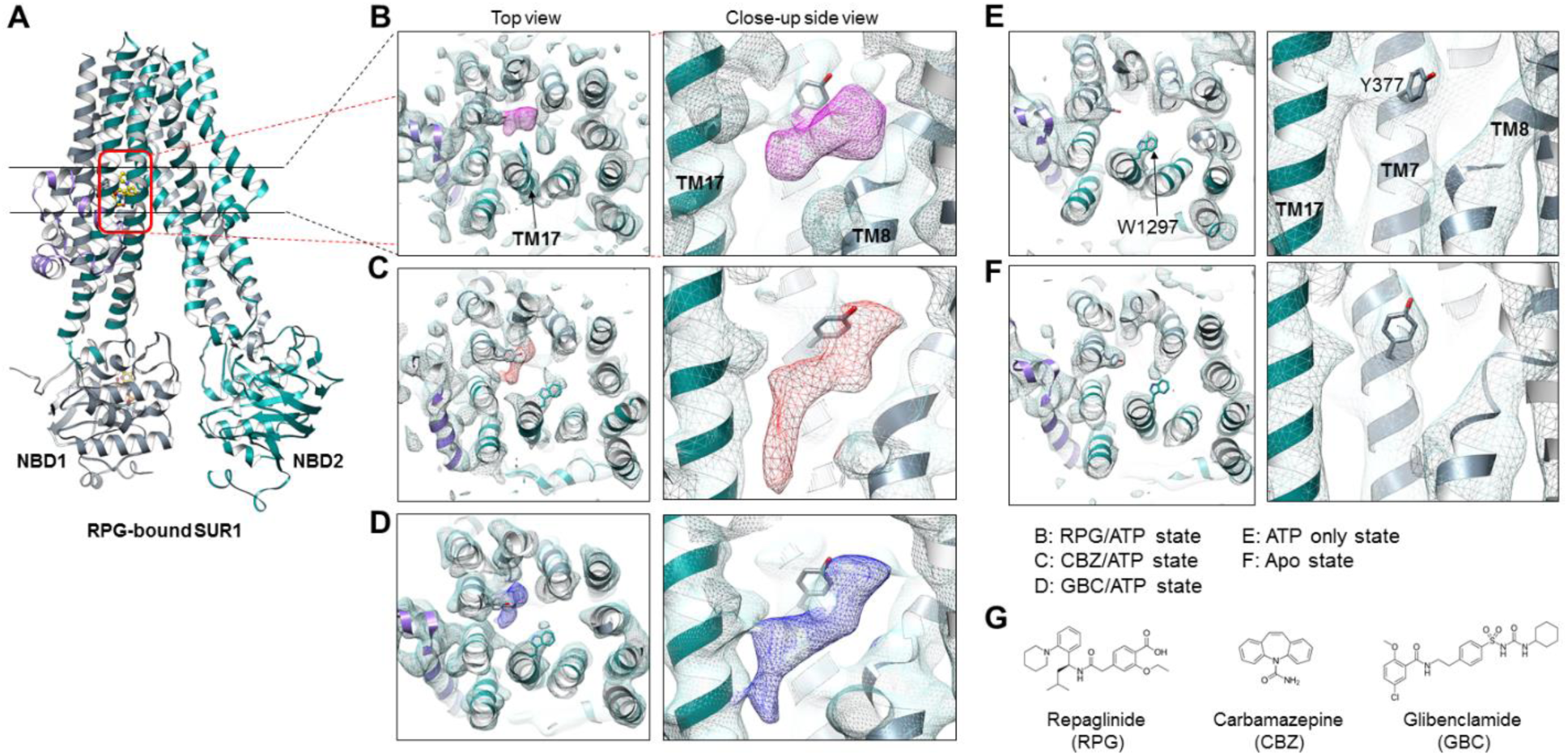
Structural comparison of the pharmacochaperone binding pocket. **(A)** Structural model of the RPG-bound SUR1 ABC transporter core module viewed from the side showing the slice viewed from the top (indicated by the two black lines) and the pocket viewed from the side at higher magnification (indicated by the red box) in B-F. **(B-F)** The pharmacochaperone pocket viewed from the top and the side of the channel in the states indicated. To enable comparison, each map was sharpened and filtered to 4.6Å (the resolution of the apo state reconstruction) with the Postprocessing procedure in RELION. Ligand density corresponding to RPG in **(B)** is shown in magenta, CBZ in **(C)** in red, and GBC in **(D)** in blue. The binding pocket is empty in both the ATP only state **(E)** and the apo state **(F)**. Note the side chain of W1297 in TM17 and Y377 in TM7 are shown and labeled in panel **(E)** to serve as reference points. **(G)** Chemical structures of the three pharmacochaperones shown in **B-D**.

Density for RPG appears compact and palm-shaped, and suggests that the molecule adopts a considerably folded shape upon binding to SUR1 (Fig. 2B; Fig.2-figure supplement 1B). Interestingly, RPG possesses a carboxyl group adjacent a benzyl group, analogous to the sulfonyl group in GBC that is also adjacent a benzyl group (Fig.2G). Refinement of an RPG molecule into the observed binding pocket density orients this carboxylate towards N1245, R1246, and R1300 (Fig.3A), which coordinate the sulfonyl group in the GBC-bound structure (Fig.3B). However, unlike GBC, the RPG density is distant from S1238. This explains previous functional data that an SUR1 S1238Y mutation does not affect RPG’s ability to modulate channel function (Hansen et al., 2002; Yan et al., 2006). The helix on the opposite side of the binding pocket (i.e. TM8; Fig.3A) is lined with hydrophobic residues (W430, F433, and L434), which may support binding through a combination of van der Waals interactions and shape complementarity. Interestingly, although similar in overall structures, we noted subtle rearrangements within the binding pocket between GBC-and RPG-bound states (Fig.3A and B). The most obvious is in W1297, which in RPG is flipped down towards the ligand (Fig.3-figure supplement 1A). Thus, there is sufficient flexibility of the binding pocket to accommodate diverse compounds with high affinity.

CBZ is a smaller molecule with molecular weight about half of that of GBC and RPG (Fig.2G). However, in the CBZ-bound SUR1 structure the cryoEM density corresponding to the ligand has a size and shape that markedly resembles GBC (Fig.2C and D), with one end pointing towards S1238. While the close proximity to S1238 is in agreement with our previous finding that mutation of S1238 to Y diminishes the ability of CBZ to both inhibit and chaperone the channel (Devaraneni et al., 2015), the density is too large to be fitted by a single CBZ molecule (Fig.2C; Fig.2-figure supplement 1C). The structure of CBZ has been extensively studied and multiple polymorphic crystalline forms have been reported, including dimers (Florence et al., 2006; Grzesiak et al., 2003). Thus, an intriguing possibility is that CBZ may bind as a dimer to occupy the entire binding pocket. Alternatively, the result could be explained by a CBZ molecule having multiple occupancies, and what is observed in the reconstruction is an average of the ensemble. Due to this uncertainly we did not model the CBZ molecule in the structure. To our knowledge this is the first protein structure determined in complex with CBZ. Thus, there are no structures for comparison. More studies are needed to distinguish these possibilities.

Structural evidence above indicates that three pharmacochaperones occupy a common binding pocket and exert their effects by engaging overlapping but non-identical sets of residues. To seek functional evidence for our CBZ and RPG binding site model, we mutated six select SUR1 residues lining the binding pocket to Ala and monitored the effect of mutation on the ability of CBZ and RPG to inhibit channel activity in ^86^Rb^+^ efflux essays, as we have done previously for GBC (Martin et al., 2017a). All six mutations reduced the sensitivity of the channel to both RPG and CBZ, but to variable extents (Fig.3C and D). For CBZ, the profile of sensitivity to the six mutations mirrors the profile previously reported for GBC (Martin et al., 2017a), consistent with the similarities in the cryoEM density for the two ligands (Fig.2C and D). By contrast, the sensitivity profile for RPG is distinct from GBC and CBZ (Fig.2D), indicating that despite occupying a common pocket, RPG engages overlapping but non-identical residues for binding. That particular mutations have distinct effects also indicate that their reduced drug sensitivities are unlikely due to indirect allosteric effects on channel structure. Among the six residues examined, Y377 is highly important for drug sensitivity for all three ligands, which agrees with our models that this residue is in close contact with the density for all three ligands. In addition, the activity of all three drugs depend similarly on interactions with R306 and N437, although to a lesser extent than Y377. By contrast, T1242 is critical for CBZ and GBC inhibition but less so for RPG inhibition, while R1300 has a greater role for RPG than the other two drugs. Of note, the modeled structure indicates that one CBZ molecule would not be able to interact simultaneously with T1242 and Y377, supporting the hypothesis that CBZ binds to SUR1 as a dimer, or adopts more than one orientation in the pocket.

**Figure 3:**
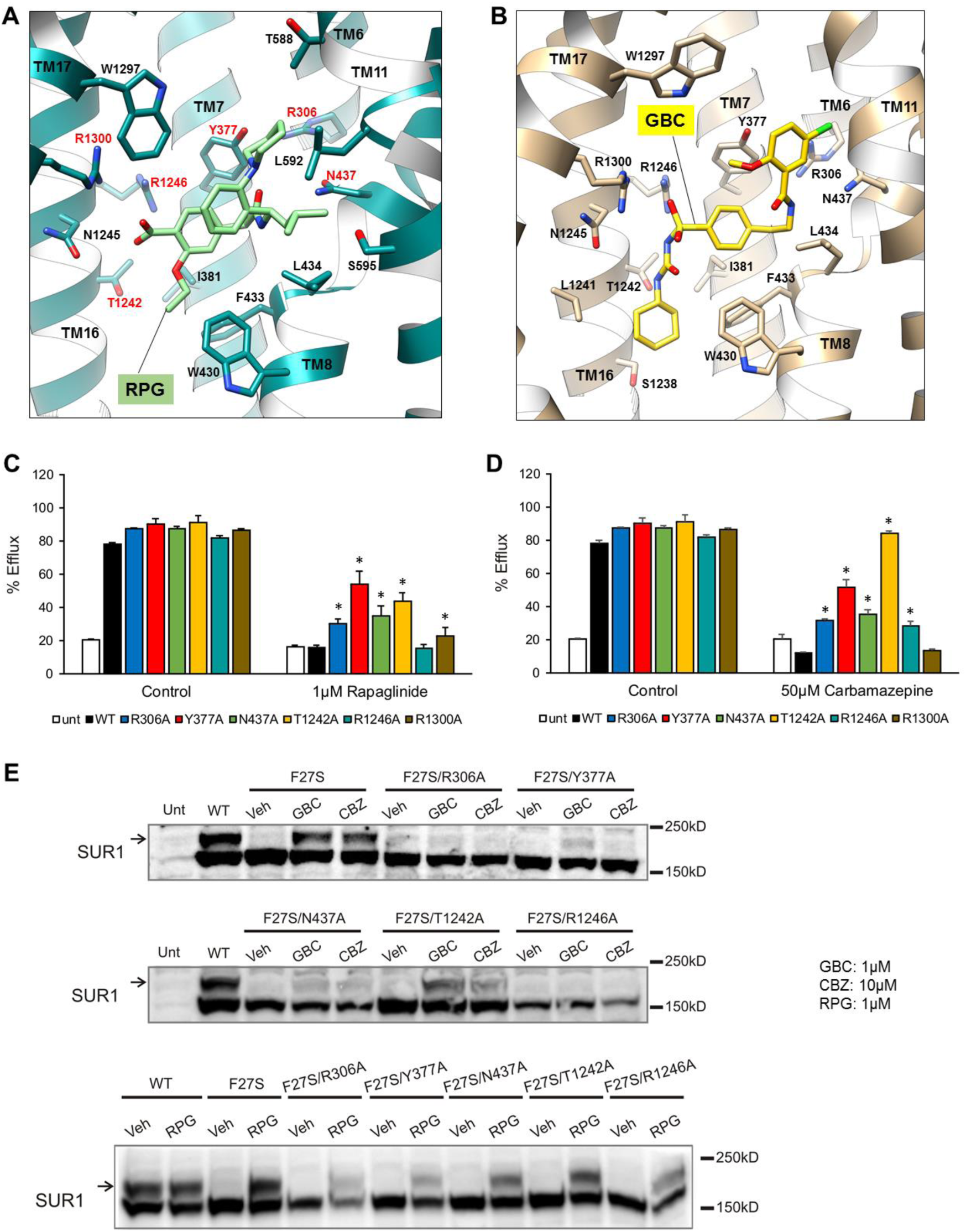
Models of the PC binding pocket. **(A)** RPG binding site model, with residues mutated to alanine in **C-E** labeled in red. **(B)** GBC binding site model. **(C-D)** Histograms of ^86^Rb^+^ efflux results showing effects of alanine mutation of select residues surrounding the bound PC on channel response to 1µM RPG (C) and 50µM CBZ (D). Asterisks indicate values significantly higher than that observed in WT in drug-treated group using paired Student’s *t*-test (*p*<0.05). Error bars represent the s.e.m. of 3-4 independent experiments. (E) Western blots showing effects of alanine mutation of the selected residues on the ability of GBC, CBZ, and RPG to correct the processing defect caused by the F27S mutation in the TMD0 of SUR1.

Next we tested the effects of the above binding pocket mutations on the ability of the three pharmacochaperones to correct the trafficking defects in SUR1-TMD0 mutants. The trafficking mutation F27S in SUR1 has been well-characterized in previous work, showing nearly undetectable mature complex-glycosylated form in the absence of pharmacochaperones but strong recoveries with both CBZ and GBC (Chen et al., 2013b). In an F27S background, we introduced five of the aforementioned six binding pocket mutations which have been shown previously to not significantly alter SUR1 maturation by themselves (Martin et al., 2017a), and examined the impact on the chaperone function of CBZ, GBC and RPG. In Western blots, there was a complete absence of the mature upper band for F27S-SUR1 in vehicle (DMSO) treated control. In contrast, there were strong upper bands when the mutant was expressed in the presence of CBZ, RPG or GBC, with signal intensities comparable to WT (Fig.3E). This chaperone ability of the drugs was attenuated or abolished for each of the binding site mutations tested in the F27S background (Fig.3E). Furthermore, mutation of W1297, which is in close proximity to GBC and RPG (Fig.3-figure supplement 1A), to alanine markedly reduced the ability of both GBC and RPG to inhibit channel activity in inside-out patch-clamp recording experiments (Fig.3-figure supplement 1B), and to rescue the processing defects of the F27S TMD0 mutation (Fig.3-figure supplement 1C). These combined functional and structural data provide strong evidence that CBZ, RPG, and GBC exert their pharmacochaperoning and channel inhibition effects by binding to a common binding pocket we have here identified.

### Structure of the distal N-terminus of Kir6.2

The N-terminal ∼30 amino acids of Kir6.2 (referred to as KNt hereinafter) has been known to be critical for channel assembly, gating, and interaction with sulfonylureas and glinides. However, underlying mechanisms remain poorly understood. Early studies showed that deletion of KNt significantly increases channel open probability (Babenko et al., 1999; Koster et al., 1999b; Reimann et al., 1999; Shyng et al., 1997), reduces ATP inhibition, and renders the channel less sensitive to sulfonylureas (Koster et al., 1999a; Reimann et al., 1999). KNt also appears to contribute to GBC binding (Kuhner et al., 2012; Vila-Carriles et al., 2007), and is necessary for high affinity interaction with RPG (Hansen et al., 2005; Kuhner et al., 2012). Deletion of amino acids 2-5 from the KNt shifts the binding affinity of RPG by more than 30-fold (Kuhner et al., 2012). Moreover, KNt is critical for pharmacochaperoning (Devaraneni et al., 2015; Schwappach et al., 2000). Removal of KNt markedly reduces the ability of GBC and CBZ to rescue SUR1-TMD0 trafficking mutations (Devaraneni et al., 2015). Our recent studies showing that *p*-azidophenylalanine genetically incorporated at Kir6.2 amino acid position 12 or 18 was photocrosslinked to SUR1 and that the extent of crosslinking increased in the presence of GBC or CBZ further suggest physical interactions between KNt and SUR1 in a drug-sensitive manner (Devaraneni et al., 2015). Taken together, these studies led us to hypothesize that KNt is located near the pharmacochaperone binding pocket we have identified.

Close examination of the three pharmacochaperone-bound SUR1 structures reconstructed using focused refinement indeed revealed significant and continuous cryo-EM density immediately adjacent to the drug binding site, especially in the RPG-bound map (Fig.4A and B). This density appears as a roughly linear peptide inserting between the two TMDs of SUR1 from the intracellular side. Interestingly, in the RPG-bound structure the piperidino moiety of RPG is in close proximity to the density that corresponds to the most N-terminus of the KNt (Fig.4-figure supplement 1A). This observation agrees with previous findings that the increased affinity to RPG caused by Kir6.2 is due to the peperidino group (Stephan et al., 2006), and that deletion of only a few amino acids from the Kir6.2 N-terminus decreases RPG binding affinity by more than 30-fold (Kuhner et al., 2012).

Of note, the KNt density is also seen in the ATP-only reconstruction though not as strong or well-defined as in the drug-bound reconstructions (i.e., the density disappears at lower σ values for the ATP-only map), but is largely absent in the apo state structure (Fig.4C). This indicates KNt enters into the central cavity of the inward-facing SUR1 even in the absence of drugs, and that drug binding stabilizes the location of KNt in the central cavity. We found that deletion of amino acids 2-5 or 2-10 markedly reduced the ability of the pharmacochaperones to correct the processing defect of the F27S TMD0 mutant SUR1 (Fig.4-figure supplement 1B), and also reduced the maturation of WT SUR1 (Fig.4-figure supplement 1C). These structure and functional observations suggest that KNt binds to the central cavity of the SUR1 ABC core structure and stabilizes SUR1-Kir6.2 association by simultaneous interaction with multiple SUR1 TMD helices, which likely enhance complex assembly during channel biogenesis and pharmacochaperoning.

### Physical and functional interactions of the distal N-terminus of Kir6.2 with SUR1

While our assignment of the KNt density is consistent with the many functional experiments described above (Babenko and Bryan, 2002; Babenko et al., 1999; Devaraneni et al., 2015; Koster et al., 1999b; Martin et al., 2016), direct physical evidence linking KNt to SUR1 residues lining the central cavity is still lacking. Identification of crosslinked SUR1 residue via *p*-azidophenylalanine engineered in KNt is technically challenging due to the complex chemistry of the azido-mediated crosslinks (Devaraneni et al., 2015). As an alternative approach, we first performed crosslinking experiments using the amine-reactive homobifunctional crosslinker DSP (dithiobis(sucinimidyl propionate)) with purified K_ATP_ channels bound to GBC, followed by mass spectrometry to identify crosslinked peptides (Nili et al., 2012). One of the inter-SUR1-Kir6.2 crosslinks we identified connected Kir6.2 lysine 5 and SUR1 lysine 602 (Fig.4-figure supplement 2), which is near the drug binding site in our structural model. The distance between Cα carbons of crosslinked lysines in our model is ∼18Å, which is within the reported range for DSP (Merkley et al., 2014). This result provides the first evidence for the physical proximity of KNt to the SUR1 ABC core central cavity.

To further corroborate the DSP crosslinking results and focus on the interaction, we attempted to crosslink KNt with SUR1 by engineered cysteine pairs. Inspection of our structural model pointed to an endogenous cysteine C1142 in SUR1 with proximity to the distal end of KNt (Fig.4-figure supplement 1A). Accordingly, we mutated Kir6.2 amino acids 2-5 to cysteine and asked whether the engineered Kir6.2 cysteines formed disulfide bonds with SUR1-C1142. We hypothesized that crosslinking of KNt with SUR1-C1142 would trap KNt within the SUR1 ABC core and reduce channel open probability, based on previous studies that deletion of KNt increases channel open probability (Babenko and Bryan, 2002; Babenko et al., 1999; Shyng et al., 1997). Using inside-out patch-clamp recording, we monitored channel activity in response to the oxidizing reagent I_2_ or the reducing reagent dithiothreitol (DTT) (Anishkin et al., 2008). Strikingly, co-expression of Kir6.2-L2C with WT SUR1 resulted in channels that displayed an accelerated current decrease in I_2_ (250µM) that was partially reversible by subsequent exposure to DTT (10mM); the other cysteine mutations did not show evidence of crosslinking with SUR1-C1142 (Fig.5). WT channels or channels formed by co-expressing Kir6.2-L2C and a SUR1 in which C1142 was mutated to an alanine (C1142A) failed to show I_2_-induced current decay that was reversed by DTT. These results offer strong validation to our structural model and support the idea that insertion of KNt into the central cavity of SUR1 ABC core serves as a mechanism to reduce channel open probability, in addition to promoting channel assembly.

**Figure 4:**
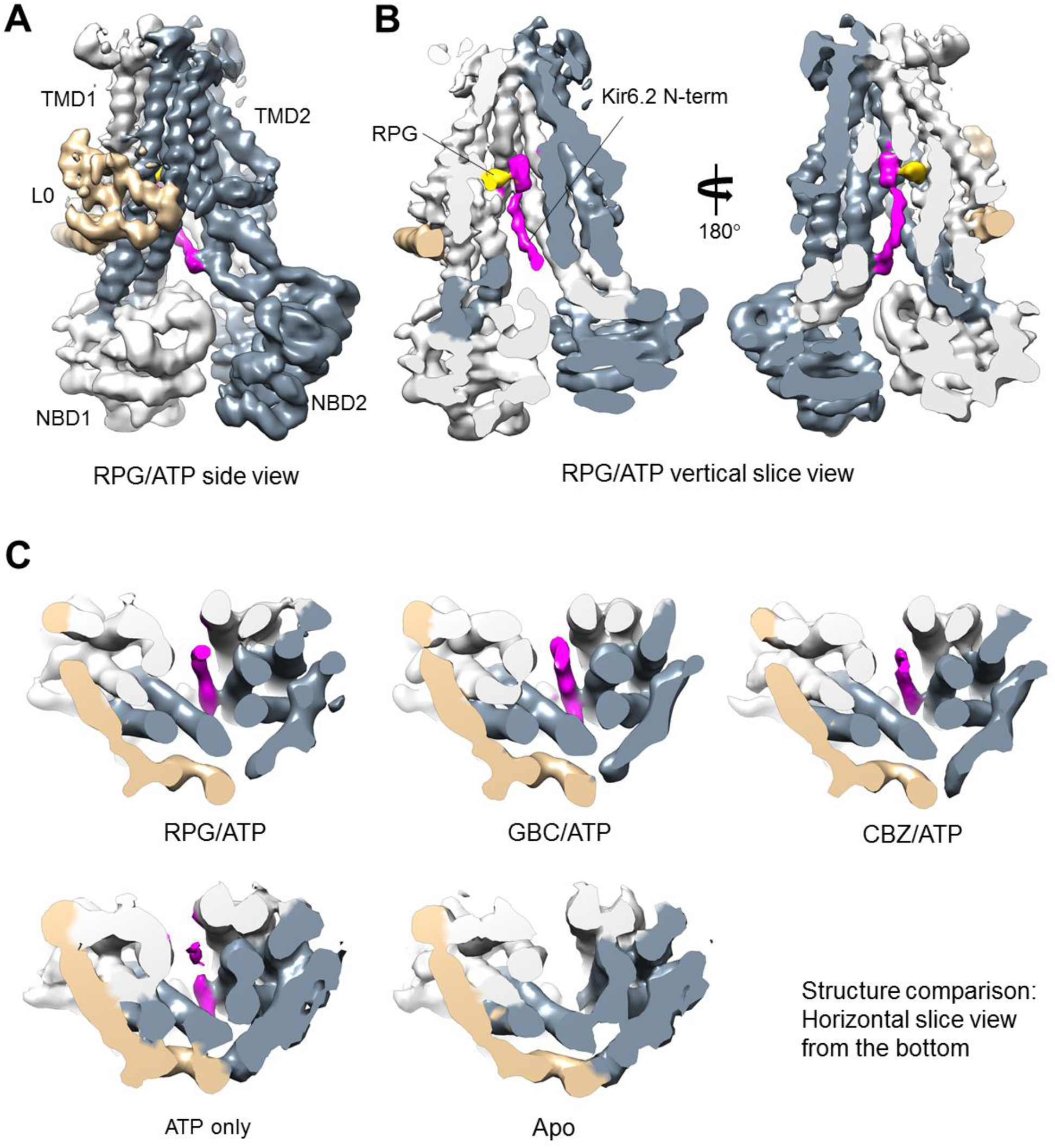
Kir6.2 N-terminus cryoEM density in SUR1. **(A)** RPG-bound SUR1 from focus-refined, unsharpened map viewed from the side. The major domains are labeled in different colors. **(B)** Vertical slide view of the map shown in **(A)** that reveals the bound RPG (in gold) and the cryoEM density of Kir6.2 N-term (in magenta). **(C)** Comparison of the Kir6.2 N-term cryoEM density in the different structures in horizontal slices viewed from the bottom. All maps are sharpened and filtered to 6Å. Apo and ATP-only structures are displayed at 1.8σ; RPG/ATP, GBC/ATP, and CBZ/ATP structures are displayed at 2.2σ. Note lower threshold is needed for the Kir6.2 N-term density in the ATP-only structure to become visible.

**Figure 5.**
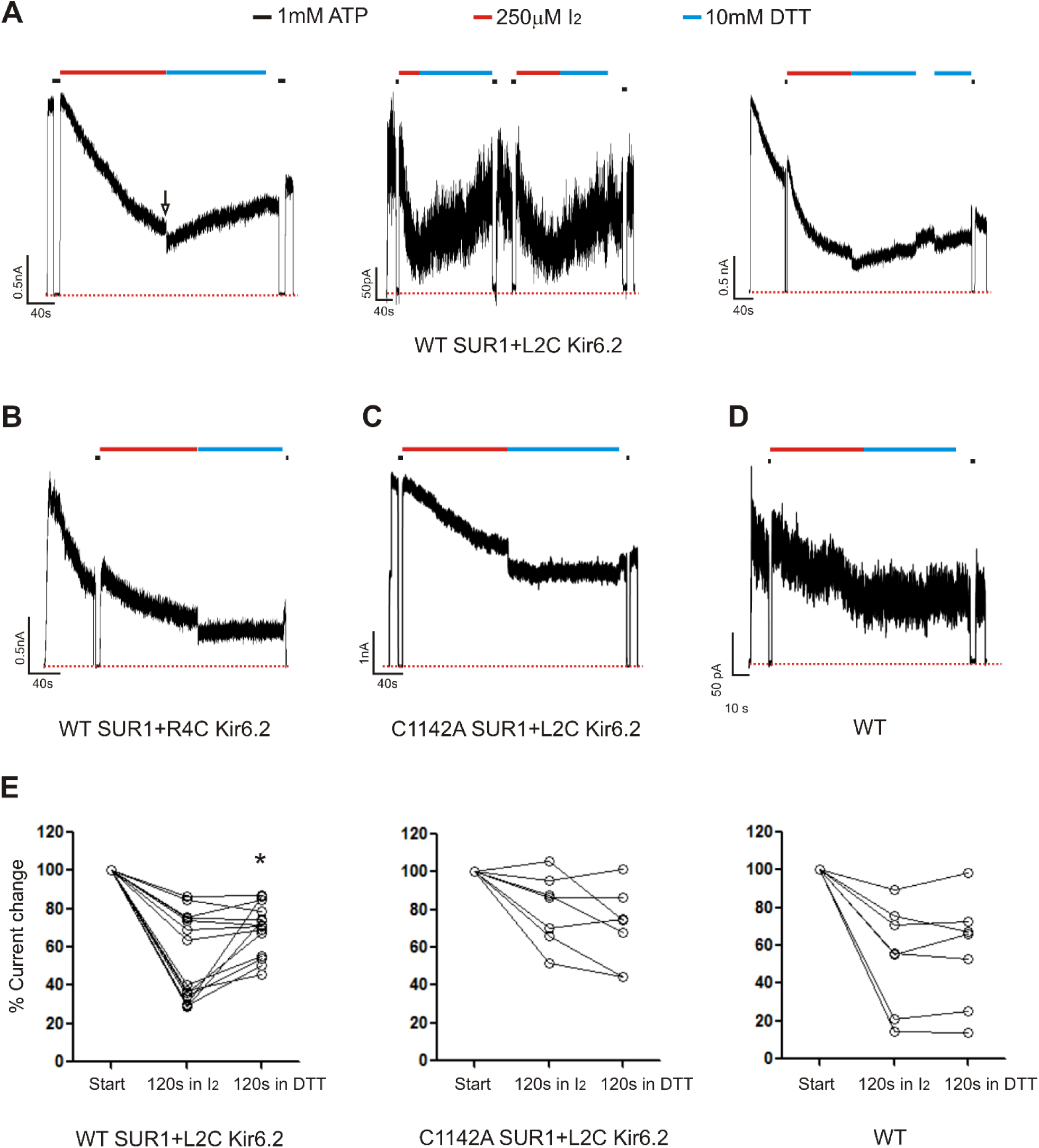
Patch-clamp recording of cysteine mutants to validate location of the Kir6.2 N-terminus. **(A)** Three select traces of channels formed by WT(C1142) SUR1 and L2C Kir6.2. Channels were exposed to 250µM I_2_ or 10mM DTT as indicated by the bars above the traces. Baseline was obtained by exposing channels to 1mM ATP. The arrow in the first trace indicates the blocking effect of the DTT that was readily reversed upon return to K-INT. **(B, C, D)** Representative traces of control channels formed by WT SUR1 and R4C Kir6.2, C1142A SUR1 and L2C Kir6.2, or WT SUR1 and Kir6.2. **(E)** Quantification of current changes at the end of 120s exposure to I_2_, and at the end of subsequent 120s exposure to DTT (expressed as % of currents at the start of I_2_ exposure). The asterisk indicates statistical significance comparing currents at the end of I_2_ exposure and the end of DTT exposure using paired student’s *t*-test (*p*<0.05).

## Discussion

Loss of membrane protein expression due to impaired folding and trafficking underlies numerous human diseases. In some cases, such defects can be overcome by small molecule ligands that bind to mutant proteins, referred to as pharmacochaperones (Bernier et al., 2004; Hanrahan et al., 2013; Powers et al., 2009; Ringe and Petsko, 2009). Although significant progress has been made in discovering pharmacochaperones, as exemplified for the CFTR mutation ΔF508 (Mijnders et al., 2017; Pedemonte et al., 2005; Veit et al., 2018), detailed structural mechanisms of how these compounds work are in most cases not well understood. This problem is even more complex in hetero-multimeric proteins such as the K_ATP_ channel where not only folding of the mutant protein itself but interactions with assembly partners may be involved. In this study, we set out to understand how structurally diverse compounds that inhibit K_ATP_ channels act as pharmacochaperones to correct channel trafficking defects. We present structural, biochemical, and functional evidence that GBC, RPG, and CBZ, despite diverse chemical structures, occupy a common pocket in SUR1, that this occupancy stabilizes a labile binding interaction between the N-terminus of Kir6.2 and the central cavity of the SUR1 ABC core, and that this interaction controls channel gating as well as the assembly and trafficking of nascent channels to the β-cell surface. Our findings shed light on the principles that govern the assembly of the K_ATP_ complex and offer insight into the mechanism by which channel inhibitors act allosterically to rescue channel trafficking defects caused by SUR1 mutations in TMD0 (see Fig.6).

**Figure 6.**
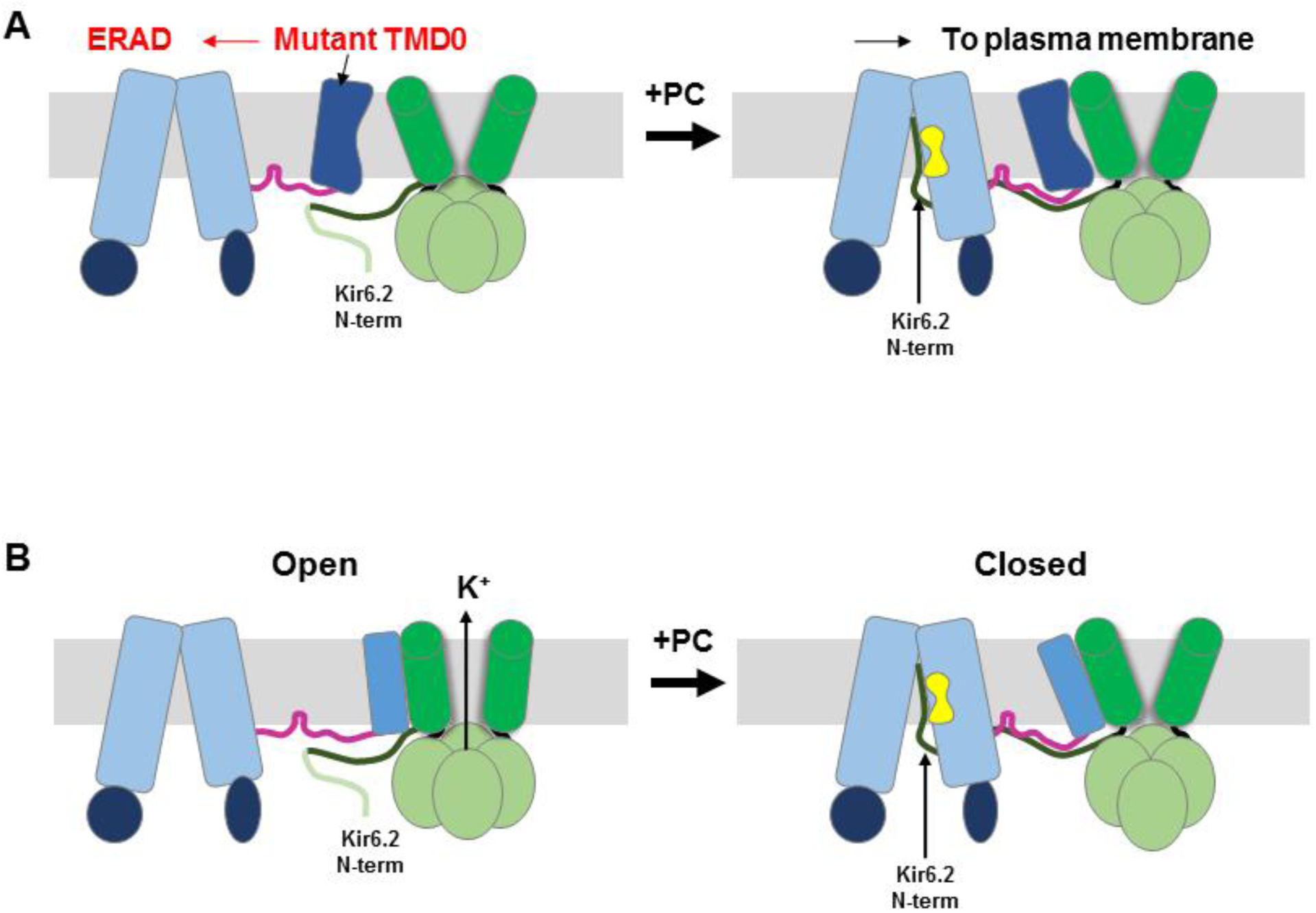
Cartoon of pharmacochaperoning and channel inhibition mechanism. **(A)** In the absence of a pharmacochaperone (PC), SUR1 containing TMD0 trafficking mutations is unable to assemble with Kir6.2 efficiently and is targeted for ER-associated degradation (ERAD). Upon binding of PC to the binding pocket in SUR1, the Kir6.2 N-term becomes stabilized in the central cavity of the SUR1 ABC core, allowing assembly of the mutant TMD0 with Kir6.2 and trafficking of the complex to the plasma membrane. **(B)** All K_ATP_ channel PCs identified to date also inhibit channel activity. In the absence of a PC, channels stay open most of the time (channel *P*_*o*_ in K-INT solution is ∼0.6-0.8). The Kir6.2 N-term is flexible and can reach into the central cavity of the SUR1 ABC core. When this happens, the channel cannot adopt a PIP_2_-bound open conformation and remains closed. Binding of PC in the SUR1 pocket stabilizes Kir6.2 N-term in the central cavity and stabilizes the channel in a closed state. In addition, PC binding stabilizes SUR1 in an inward-facing conformation, unable to be stimulated by Mg-nucleotides. Crosslinking of SUR1’s endogenous C1142 with engineered Kir6.2 L2C also traps the Kir6.2 N-term in the central cavity, closing the channel (see **Fig.5**).

Among ABC transporters and inward rectifier potassium channels, SUR1 and Kir6.2 are unique in having evolved to become functionally inter-dependent. The mechanisms by which these two structurally unrelated proteins form a heteromeric complex to regulate K^+^ transport and insulin secretion have been intensively investigated. Recently, several cryo-EM structures have been reported of K_ATP_ channels naturally assembled from separate SUR1 and Kir6.2 proteins (Li et al., 2017; Martin et al., 2017a; Martin et al., 2017b), or comprising engineered SUR1(C)-(N)Kir6.2 fusion proteins (Lee et al., 2017; Wu et al., 2018). These structures reveal the channel’s overall architecture as a Kir6.2 tetramer surrounded by four SUR1, wherein Kir6.2 makes direct contact with the TMD0 of SUR1. This structure explains why numerous trafficking mutations are found in TMD0 (Snider et al., 2013). However, we have shown in the structure of GBC-bound K_ATP_ complexes that, GBC resides well within the TM bundle of the SUR1 ABC core, above NBD1 and distant from TMD0 (Martin et al., 2017a). An allosteric mechanism by which binding of GBC rescues TMD0 trafficking mutations has remained obscure.

A critical missing piece of the puzzle was the KNt, which we have shown to be involved in pharmacochaperone rescue of TMD0 mutations (Devaraneni et al., 2015). KNt has not been resolved in previously determined structures (Li et al., 2017; Martin et al., 2017a; Martin et al., 2017b), although weak and disconnected cryoEM densities in the central cavity of the SUR1 ABC core have fueled speculation (Sikimic et al., 2018; Wu et al., 2018). Here, through improved focused refinement algorithms, cross-correlation of multiple structures, and biochemical as well as functional crosslinking experiments, we present evidence necessary to properly assign the KNt density adjacent to the pharmacochaperone binding site within the central cavity of the SUR1 ABC core.

The combined structural and functional studies presented here provide a clear mechanism by which pharmacochaperone binding acts in *trans* to overcome assembly defects caused by mutations within TMD0. Because the pharmacochaperone binding site is formed by the TM helices of the SUR1 ABC core, and TMD0 is a separate domain, it is unlikely that its occupancy by chemically diverse compounds overcome or prevent misfolding of TMD0 trafficking mutations at the site of the mutation. Supporting this notion, GBC, RPG, and CBZ fail to chaperone TMD0 trafficking mutants in the SUR1_RKR-AAA_ background, a variant in which the RKR ER retention signal has been mutated to AAA to allow Kir6.2-independent trafficking to the cell surface (Devaraneni et al., 2015; Yan et al., 2006). Rather, occupancy of the pharmacochaperone binding site stabilizes the insertion of KNt into the central cavity of the SUR1 ABC core, an otherwise labile interaction that is nonetheless crucial for assembly and trafficking of the channel complex out of the ER (Fig.6A). Insertion of KNt into the central cavity of the SUR1-ABC core as a principal mechanism to stabilize SUR1-Kir6.2 interaction during assembly is consistent with previous findings (Devaraneni et al., 2015) and our current results (Fig.4-figure supplement 1C) that deletion of KNt greatly reduced the biogenesis efficiency of WT channels.

The location of the KNt density observed in our structures, including the ATP-only structure, also illuminates how Kir6.2 and SUR1 interact to modulate channel activity. First, by occupying the central cavity of the SUR1 ABC core, KNt prevents NBD dimerization and stabilizes SUR1 in an inward-facing conformation, thereby abolishing the ability of Mg-nucleotides to stimulate channel activity. Second, by trapping KNt in the central cavity, SUR1 likely prevents the Kir6.2 tetramer from adopting a PIP_2_-bound open conformation. Our finding that crosslinking of SUR1-C1142 with Kir6.2-L2C accelerates current decay that is reversible by DTT lends support to the second mechanism. It also explains early studies showing that deletion of KNt increased channel open probability and decreased the ability of sulfonylureas to inhibit the channel (Babenko and Bryan, 2002; Koster et al., 1999a; Reimann et al., 1999; Shyng et al., 1997). That KNt EM density, albeit weaker, is found in the ATP-only structure suggests KNt spends significant time in SUR1’s ABC core cavity when Kir6.2 is in the ATP-bound closed state. By contrast, the KNt cryoEM density is absent in the apo structure, suggesting that when no ATP is bound to Kir6.2, KNt has little residence time in the ABC core central cavity. The fact that despite not seeing KNt, the Kir6.2 channel is still closed in the dominant class structure is likely due to loss of endogenous PIP_2_ during purification.

Our structures show that CBZ and RPG share the same binding pocket as GBC in K_ATP_ channels. Although both RPG and CBZ have been characterized extensively, with RPG being a commonly used oral hypoglycemic medication and CBZ an anticonvulsant, there are no known structures of these drugs bound to proteins. Thus, our structures provide the first examples of how these drugs interact with their target proteins. Despite occupying a common pocket, each of the three compounds appear to engage a unique set of residues. This is particularly clear when comparing GBC and RGP. Curiously, we found that the density corresponding to CBZ cannot be fit with a single molecule. CBZ is a well-known Na_v_ channel blocker (Lipkind and Fozzard, 2010). It will be important to determine whether CBZ adopts a similarly bound arrangement in other target proteins.

Despite functional divergence, all ABC transporters share some structural similarities, in particular the ABC core structure (Thomas and Tampe, 2018). An exciting finding here is how the Kir6.2 N-terminus can act as a plug or handle to regulate channel gating and assembly by inserting itself into the ABC core central cavity formed by the two TM bundles. The exploitation of this structural space is akin to the mechanism by which viral peptide ICP47 enables immune-evasion of the pathogens (Blees et al., 2017; Oldham et al., 2016; Parcej and Tampe, 2010). In this case, ICP47 secreted by viruses such as Herpes simplex virus and Cytomegalovirus inserts into the inner vestibule formed by the ABC peptide transporters TAP-1 and TAP-2 (transporter associated with antigen processing) (Blees et al., 2017; Oldham et al., 2016), thus preventing the transport of cytosolic peptides into the ER for MHC complex loading and immune surveillance. Following this, molecules which can reside within this space may offer opportunities for functional modulation of ABC transporters such as K_ATP_. Indeed, application of a Kir6.2 N-terminal peptide (a.a. 2-33) to the cytoplasmic face of K_ATP_ channels in isolated membrane patches has been shown to increase K_ATP_ channel open probability (Babenko and Bryan, 2002). This is presumably due to competition of the exogenous peptide with the N-terminus of Kir6.2 for binding to the SUR1 ABC core. With regard to the drug binding pocket, it is worth noting that both GBC and CBZ have been reported to bind or are substrates of other ABC transporters (Zhou et al., 2008). For example, Bessadok et al. (Bessadok et al., 2011) recently showed that the multidrug resistance transporter P-glycoprotein (ABCB1) recognizes several SUR1 ligands including GBC, albeit with much lower affinity. Moreover, CBZ has been shown to correct the trafficking defect of ΔF508 CFTR (ABCC9), the most prevalent mutation underlying cystic fibrosis (Carlile et al., 2012). An intriguing possibility is that these other ABC transporters recognize SUR1 ligands through a similar binding pocket identified here via conserved residues within the SUR1 GBC binding pocket such as R1246 and W1297. It would be interesting to determine in the future whether and how CBZ binding affects ΔF508 CFTR structure to correct its processing defect (Yan et al., 2006).

In summary, we have uncovered mechanisms by which a diverse set of compounds modulate the gating and assembly/trafficking of the pancreatic K_ATP_ channel, a critical regulator of glucose-stimulated insulin secretion. Our findings may serve as a drug development platform in K_ATP_ channels and other ABC transporters for new pharmacological chaperones or modulators with improved efficacy and specificity.

## Materials and Methods

### Protein expression and purification

K_ATP_ channels were expressed and purified as described previously (Martin et al., 2017a; Martin et al., 2017b). Briefly, the genes encoding pancreatic K_ATP_ channel subunits, which comprise a hamster SUR1 and a rat Kir6.2 (94.5% and 96.2% sequence identity to human, respectively), were packaged into recombinant adenoviruses (Lin et al., 2005; Pratt et al., 2009); both are WT sequences, except for a FLAG tag (DYKDDDDK) engineered into the N-terminus of SUR1 for affinity purification. INS-1 cells clone 832/13 (Hohmeier et al., 2000), a rat insulinoma cell line, were infected with the adenoviral constructs in 15 cm tissue culture plates. Protein was expressed in the presence of 1mM Na butyrate as well as 5µM RPG (for the RPG/ATP structure) or 10µM CBZ (for the CBZ/ATP structure) to enhance expression and formation of the channel complex (Chen et al., 2013b; Yan et al., 2006). At 40-48 hours post-infection, cells were harvested and cell pellets flash frozen in liquid nitrogen and stored at −80°C until purification.

For purification, cells were resuspended in hypotonic buffer (15mM KCl, 10mM HEPES, 0.25 mM DTT, pH 7.5) and lysed by Dounce homogenization. The total membrane fraction was resuspended in buffer A (0.2M NaCl, 0.1M KCl, 0.05M HEPES, 0.25mM DTT, 4% sucrose, pH 7.5) and solubilized with 0.5% Digitonin. Note for the ATP-only dataset, 1mM ATP was included throughout the purification; for the CBZ/ATP dataset, 10µM CBZ and 1mM ATP were present throughout; for the RPG/ATP dataset, 1mM ATP and 30µM RPG were included throughout. The soluble fraction was incubated with anti-FLAG M2 affinity agarose for 4 hours and eluted with buffer A (without sucrose) containing 0.25 mg/mL FLAG peptide. Purified complexes were concentrated to ∼1-1.5 mg/mL and used immediately for cryo grid preparation. For the RPG/ATP sample, the final concentration of RPG was 30µM, and ATP 1mM; for the CBZ/ATP sample, the final concentration of CBZ was 10µM, and ATP 1mM; for the ATP-only sample, the final ATP concentration was 1mM.

### Sample preparation and data acquisition for cryo-EM analysis

3 µL of purified K_ATP_ channel complex was loaded onto UltrAufoil gold grids which had been glow-discharged for 60 seconds at 15 mA with a Pelco EasyGlow ®. The sample was blotted for 2s (blot force −4; 100% humidity) and cryo-plunged into liquid ethane cooled by liquid nitrogen using a Vitrobot Mark III (FEI).

Single-particle cryo-EM data was collected on a Titan Krios 300 kV using a Falcon III detector (Thermo Scientific) for the RPG/ATP dataset and a Gatan K2 Summit detector for the CBZ/ATP and the ATP only datasets. The apo dataset was collected on a Talos Arctica 200 kV microscope with a Gatan K2 Summit detector using SerialEM. Data collected using the Gatan K2 Summit direct electron detector were performed in the super-resolution mode, post-GIF (20eV window), at a physical pixel size of 1.72Å (Krios) or 1.826Å (Arctica). For the RPG/ATP dataset, the calibrated pixel-size of the Falcon III was at 1.045Å. Defocus was varied between −1.0 and −3.0µm across the datasets. Detailed imaging parameters are provided in Table 1.

### Image processing

The raw frame stacks were gain-normalized and then aligned and dose-compensated using Motioncor2 (Zheng et al., 2017) with patch-based alignment (5×5). CTF parameters were estimated from the aligned frame sums using CTFFIND4 (Rohou and Grigorieff, 2015). Particles were picked automatically using DoGPicker (Voss et al., 2009) with a broad threshold range to reduce bias. Subsequently, each image was analyzed manually to recover particles missed by automatic picking and remove bad micrographs. 2D classifications were done using RELION-2 (Kimanius et al., 2016). Classes displaying fully assembled complexes and high signal/noise were selected and particles re-extracted at 1.72 Å/pix (or 1Å/pix for the RPG/ATP dataset) and then used as input for 3D classification in RELION-2 (see Table 1 and Fig.1-figure supplements 1-4).

Extensive 3D classification was performed to sample heterogeneity within the data. Symmetry was not imposed at this step in order to select only true four-fold classes. Up to 4 consecutive rounds of classification were performed, specifying 4 or 5 classes per round. Individual classes and combinations of classes were refined for each dataset to achieve the best reconstructions (Fig.1-figure supplements 1-4). A soft mask encompassing the entire complex was used during refinement in RELION, with C4 symmetry imposed to yield an overall channel reconstruction using the gold-standard FSC cutoff. The map was B-factor corrected and filtered using RELION-2’s Postprocessing procedure, with the same mask used for refinement.

### Focused refinement

Focused refinement of SUR1 was carried out in RELION-3 using symmetry expansion, partial signal subtraction that removes signals outside the masked region, followed by further 3D refinement of signal subtracted particles (Scheres, 2016). Different masking strategies were applied to the different datasets for focused refinement to obtain the best reconstruction (Fig.1-figure supplements 1-4). For the RPG/ATP dataset, refinement against the SUR1 ABC core module following symmetry expansion and signal subtraction led to significantly improved resolution (Fig.1-figure supplement 1). For the CBZ/ATP dataset, refinement against the entire SUR1 was performed, which also yielded an improved density map for SUR1 (Fig.1-figure supplement 2)

Similar focused refinement strategies were initially applied for the previously published GBC/ATP dataset and the new ATP-only and the apo datasets. However, this resulted in deterioration of map quality and resolution compared to the starting reconstructions. Therefore, alternative signal subtraction strategies were tested for focused refinement, which led to the use of a mask that includes the Kir6.2 tetramer and one SUR1 for the three datasets. Thus, for the GBC/ATP dataset, the particles included in the final C4 reconstruction were subjected to C4 symmetry expansion and signal subtraction using a mask that includes the Kir6.2 tetramer and one SUR1 (Fig.1-figure supplement 3). Although the resulting map showed little improvement in overall resolution compared to our previously published C4 map (Martin et al., 2017a), the local resolution of the SUR1 in several regions was significantly improved, especially NBD1 and the linker between TMD2 and NBD2 (Fig.1-figure supplement 5). A similar scheme with some modifications was employed for the ATP-only and the apo datasets (Fig.1-figure supplements 4).

### Modeling

Modeling was performed for the RPG- and GBC-bound SUR1 maps, which following focused refinement have the highest resolutions among all the structures included in this study.

For RPG-bound SUR1, the final map was B-factor corrected and filtered using RELION-3’s Postprocessing tool to optimize observable side chain features. For model building, we used SUR1 from our previously published structure (PDB:6BAA) as the initial model. The ABC core structure including the two transmembrane domains (TMD1 and TMD2) and two cytosolic nucleotide binding domains (NBD1 and NBD2) was docked into the density map by rigid-body fitting using Chimera’s “Fit in” tool. The model was further optimized by rigid-body refinement using “Phenix.real_space_refinemet” with default parameters (Afonine et al., 2018). Of note, additional densities not observed in the previously published K_ATP_ cryoEM density maps are now clearly resolved, particularly the ATP density in NBD1 and the density for the linker regions between TMD2 and NBD2 (Fig.1-figure supplements 5). To build additional residues in these regions, the final map was sharpened by B-factor using RELION 3.0 (Zivanov et al., 2018) to optimize observable map features. As there are no homology models available for these disordered regions, models were built manually de novo in COOT (Emsley et al., 2010) followed by refinement in PHENIX, as detailed below. Initially, poly-alanines were built into the maps with some clear bulky side chains in COOT; refinement was performed by “Phenix.real_space_refine” with the following parameters: global minimization, morphing, and SUR1 initial model as a restraining reference model. Bulky side chains in the new loop regions were added, and the region between D1060-C1079 that was previously modeled as a non-helical structure was corrected to a complete helix by adjustment of residues in accordance with the density map. The model was refined by iterative manual inspection and side chains adjustment to fit the densities in COOT and real space refinement using “Phenix.real_space_refine” with tight secondary structure and torsion angle restrains. Previously published NBD domains (PDB: 6BAA) was inserted as a reference model to provide additional restrain and to minimize overfitting. The Kir6.2 N-terminus (amino acids 1-19) was built as a poly-alanine model with the side chain of L2 modeled to show its relation to SUR1 C1142. For modeling RPG in the cryoEM density map, a molecular topology profile for RPG was created using eLBOW in PHENIX, and refined into the cryoEM density in COOT. The RPG molecule with new coordination was then added to the ABC core structure model for further refinement.

Model building for the GBC-SUR1 map was similarly performed. Because the GBC-bound SUR1 map was refined using the Kir6.2 tetramer with one full SUR1 subunit as a module, we created an initial model for SUR1 by merging TMD0 from the previous model (PDB:6BAA) with the final ABC core structure built using the RPG-SUR1 map described above, which was then docked into the GBC-SUR1 map using Chimera and refined using COOT and PHENIX. GBC was docked into the ligand density in the GBC-bound SUR1 map derived from focused refinement. In addition to the regions noted above for the RPG-SUR1 map, densities corresponding to K329-G353 are of sufficient quality to build poly-alanines with some bulky side chains de novo in COOT. Extra density at Asn10 likely corresponds to glycosylated Asn10, although the sugar moiety is insufficiently resolved for modeling.

SUR1 model building in the CBZ-bound SUR1 map was also performed following the same procedure outlined for GBC-SUR1, except that the initial model was the final ABC core structure from the RPG-SUR1 model. Note for the CBZ map, no attempt was made to incorporate the CBZ molecule due to uncertainty in whether the CBZ binds as a monomer or dimer, as discussed in the main text. Also, due to insufficient resolution ATP molecule bound at NBD1 was not modeled.

### Functional studies

Point mutations were introduced into hamster SUR1 cDNA in pECE using the QuikChange site-directed mutagenesis kit (Stratagene). Mutations were confirmed by DNA sequencing. Mutant SUR1 cDNAs and rat Kir6.2 in pcDNA1 were co-transfected into COS cells using FuGENE®6, as described previously (Devaraneni et al., 2015), and used for Western blotting, ^86^Rb^+^ efflux assays, and electrophysiology as described below.

For Western blotting, cells were lysed in a buffer containing 20 mM HEPES (pH 7.0), 5 mM EDTA, 150 mM NaCl, 1% Nonidet P-40, and CompleteTR protease inhibitors (Roche) 48-72 hours post-transfection. To rescue trafficking-impaired SUR1-F27S mutant, 0.1% DMSO (vehicle control), 1µM GBC, 1µM RPG or 10µM CBZ were added to cells 24 hours before cell harvest. Proteins in cell lysates were separated by SDS/PAGE (8%), transferred to nitrocellulose membrane, probed with rabbit anti-SUR1 antibodies against a C-terminal peptide of SUR1 (KDSVFASFVRADK), followed by HRP-conjugated anti-rabbit secondary antibodies (Amersham Pharmacia), and visualized by chemiluminescence (Super Signal West Femto; Pierce) with FluorChem E (ProteinSimple).

For ^86^Rb^+^ efflux assays, cells were plated and transfected in 12-well plates. 24-36 hours post-transfection, cells were incubated overnight in medium containing ^86^RbCl (0.1 μCi/ml). The next day, cells were washed in Krebs-Ringer solution twice and incubated with metabolic inhibitors (2.5μg/ml oligomycin and 1mM 2-deoxy-D-glucose) in Krebs-Ringer solution for 30 min in the presence of ^86^Rb^+^. Following two quick washes in Krebs-Ringer solutions containing metabolic inhibitors and 0.1% DMSO (vehicle control), 1µM RPG, or 50µM CBZ, 0.5 ml of the same solution was added to each well. At the end of 2.5 minutes, efflux solution was collected for scintillation counting, new solution added, and steps repeated for 5, 7.5, 15, 25, and 40 min cumulative time points. After the 40 min time point efflux solution was collected, cells were lysed in Krebs-Ringer containing 1% SDS. ^86^Rb^+^ in the solution and the cell lysate was counted. The percentage efflux was calculated as the radioactivity in the efflux solution divided by the total activity from the solution and cell lysate, as described previously (Chen et al., 2013a; Yan et al., 2007).

For electrophysiology experiments to test the effects of crosslinking, cells co-transfected with SUR1 and Kir6.2 along with the cDNA for the green fluorescent protein GFP (to facilitate identification of transfected cells) were plated onto glass coverslips 24 hours after transfection and recordings made in the following two days. All experiments were performed at room temperature as previously described (Devaraneni et al., 2015). Micropipettes were pulled from non-heparinized Kimble glass (Fisher Scientific) on a horizontal puller (Sutter Instrument, Co., Novato, CA, USA). Electrode resistance was typically 1-2 MO when filled with K-INT solution containing 140 mM KCl, 10 mM K-HEPES, 1 mM K-EGTA, pH 7.3. ATP was added as the potassium salt. Inside-out patches of cells bathed in K-INT were voltage-clamped with an Axopatch 1D amplifier (Axon Inc., Foster City, CA). ATP (as the potassium salt), I_2_ (diluted from 250mM stock in ethanol), or dithiothreitol (DTT) were added to K-INT as specified in the figure legend. All currents were measured at a membrane potential of −50 mV (pipette voltage = +50 mV). Data were analyzed using pCLAMP10 software (Axon Instrument). Off-line analysis was performed using Microsoft Excel programs. Data were presented as mean ± standard error of the mean (s.e.m).

### DSP crosslinking and mass spectrometry

For each crosslinking experiment, ∼10µg purified K_ATP_ channels were incubated with the amine-reactive, homobifunctional crosslinker DSP [dithiobis(succinimidyl propionate); 12Å arm length], at a final concentration of 1mM. The crosslinking reaction was allowed to proceed for 20min on ice, and was performed in the presence of 1 μM GBC. The reactions were quenched with 100mM Tris, pH 8.0, and crosslinked protein was precipitated with chloroform/methanol. Generation of tryptic peptides for mass spectrometry was performed with the aid of ProteaseMax (Promega) detergent, according to the manufacturer’s protocol, with the omission of reduction and alkylation steps in order to preserve the disulfide bond within DSP. Briefly, precipitated protein was resuspended directly in 0.1% ProteaseMax/50mM ammonium bicarbonate/3.4M urea for 60 min at 37°C. Trypsin was added to a final concentration of 0.18µg/ml and protein was digested for 4 hours at 37°C. Trifluoroacetic acid was added to 0.5% to inactivate trypsin and digests were incubated at room temperature for 5 min followed by centrifugation at 20,000x *g* for 10min to remove insoluble material. Digested peptides were then analyzed by an Orbitrap Fusion mass spectrometer equipped with an electron transfer dissociation (ETD) source that cleaves the disulfide bond as detailed below. In total, the experiments revealed nine SUR1-SUR1 and three SUR1-Kir6.2 crosslinks.

Peptide digests were dried by vacuum centrifugation and dissolved in 20 µl of 5% formic acid and injected at a flow rate of 5 μl/min onto an Acclaim PepMap 100 μm × 2 cm NanoViper 5-μm C18 trap (Thermo Scientific) using mobile phase A containing water and 0.1% formic acid. After 5 min, the trap was switched in-line with a PepMap RSLC C18, 2 μm, 75 μm × 25 cm EasySpray column fitted in an EasySpray nano electrospray source (Thermo Scientific) at 40°C. Peptides were eluted with a 90 min gradient of 7.5-30% mobile phase B containing acetonitrile and 0.1% formic acid at a flow rate of 0.3 μl/min. Mass spectrometry data were collected using an Thermo Scientific Orbitrap Fusion instrument to identify DSP crosslinked peptides. Briefly, survey MS scans were acquired at 120,000 resolution and peptides with charge states of +4 to +8 were selected for data-dependent MS2 scans. These scans were acquired at 30,000 resolution in the Orbitrap mass analyzer following electron transfer dissociation (ETD) to selectively fragment the disulfide bond in the DSP crosslinker, as previously described during the mapping of disulfides in native proteins (Nili et al., 2013; Nili et al., 2012; Wu et al., 2009). This was followed by data-dependent MS3 scans on the top 6 most intense ions in the MS2 scans using collision-induced dissociation in the instrument’s ion trap.

DSP crosslinked peptides liberated by EDT fragmentation were identified by Sequest searches of MS3 scans using Protein Discoverer 1.4 (Thermo Scientific). The searches were performed using a Swiss Prot rat database downloaded in May 2015 containing 8078 entries supplemented with the sequences of hamster SUR1. Searches were configured to use only MS3 scans, MS(n - 1) precursor masses, differential modifications of 15.995 Da on methionines (oxidation), and +87.998 Da on lysines (mass of DSP crosslinker with cleaved disulfide). Precursor and fragment ion tolerances were 2 and 0.6 Da, respectively, using monoisotopic mass calculations for both, and y and b ion fragments were specified in the search. Approximate peptide false discovery rates were calculated using a sequence reversed decoy database strategy and Percolator software (Kall et al., 2007). Only peptides with q values of < 0.05 were accepted. Lists of identified peptides were then exported and filtered so that only peptides derived from Kir6.1/SUR1 that also contained the expected +87.998 mass shift at lysine residues were considered. Peptides falling within 8 scan numbers of one another were further investigated to confirm that they were derived from the same parent ion during a MS2 scan, and had summed masses that were consistent with the mass of the original crosslinked peptide detected in the initiating MS1 scan.

## Acknowledgements

We thank Dr. Christopher B. Newgard for the INS-1E cell line clone 832/13 and the staff of the Multi-Scale Microscopy Core at the Oregon Health & Science University for help with imaging and data collection. We also thank Dr. Bruce Patton for comments on the manuscript. This work was supported by US National Institutes of Health grants DK57699 (S.-L. S.), DK066485 (S.-L. S.) and F31 DK105800 (G.M.M.).

## Author Contributions

GMM conceived, designed, and conducted experiments, analyzed data, and prepared the manuscript; MWS analyzed cryoEM data and prepared the manuscript; ZY, LI, and BK conducted experiments; LLD conducted the mass spectrometry experiments, made Fig.4-figure supplement 2, and edited the manuscript; CY supervised data collection and analyzed data, and edited the manuscript; SLS conceived, designed, and conducted experiments and prepared the manuscript.

---

**Figure 1S1-figure supplement 1:**
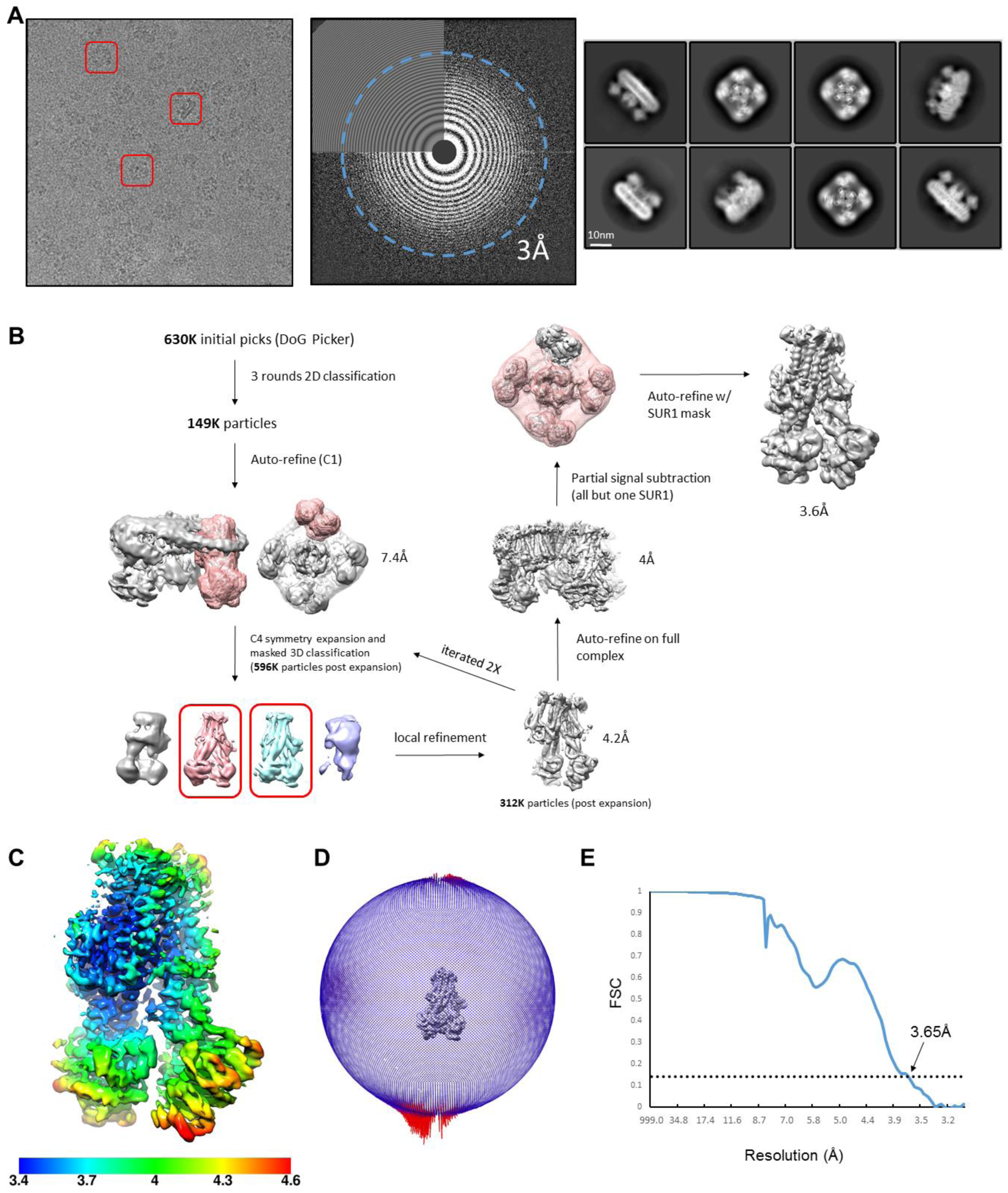
Data collection and image processing workflow for the RPG/ATP state. **(A)** Left: Representative micrograph after alignment with Motioncor2. A few K_ATP_ channel complexes of various orientation are outlined by the red box. Middle: Power spectrum calculated with Ctffind4, with resolution reaching 3.0Å. Right: Representative 2D classes. **(B)** Overview of data processing workflow. Particle picking was performed automatically with DoGPicker and manual inspection. All other image processing steps were performed in RELION-3. **(C)** Local resolution plot of focal refined SUR1 map. **(D)** Angular distribution plot. **(E)** Fourier shell correlation (FSC) of two independent half maps of focal refined SUR1.

**Figure 1-figure supplement 2:**
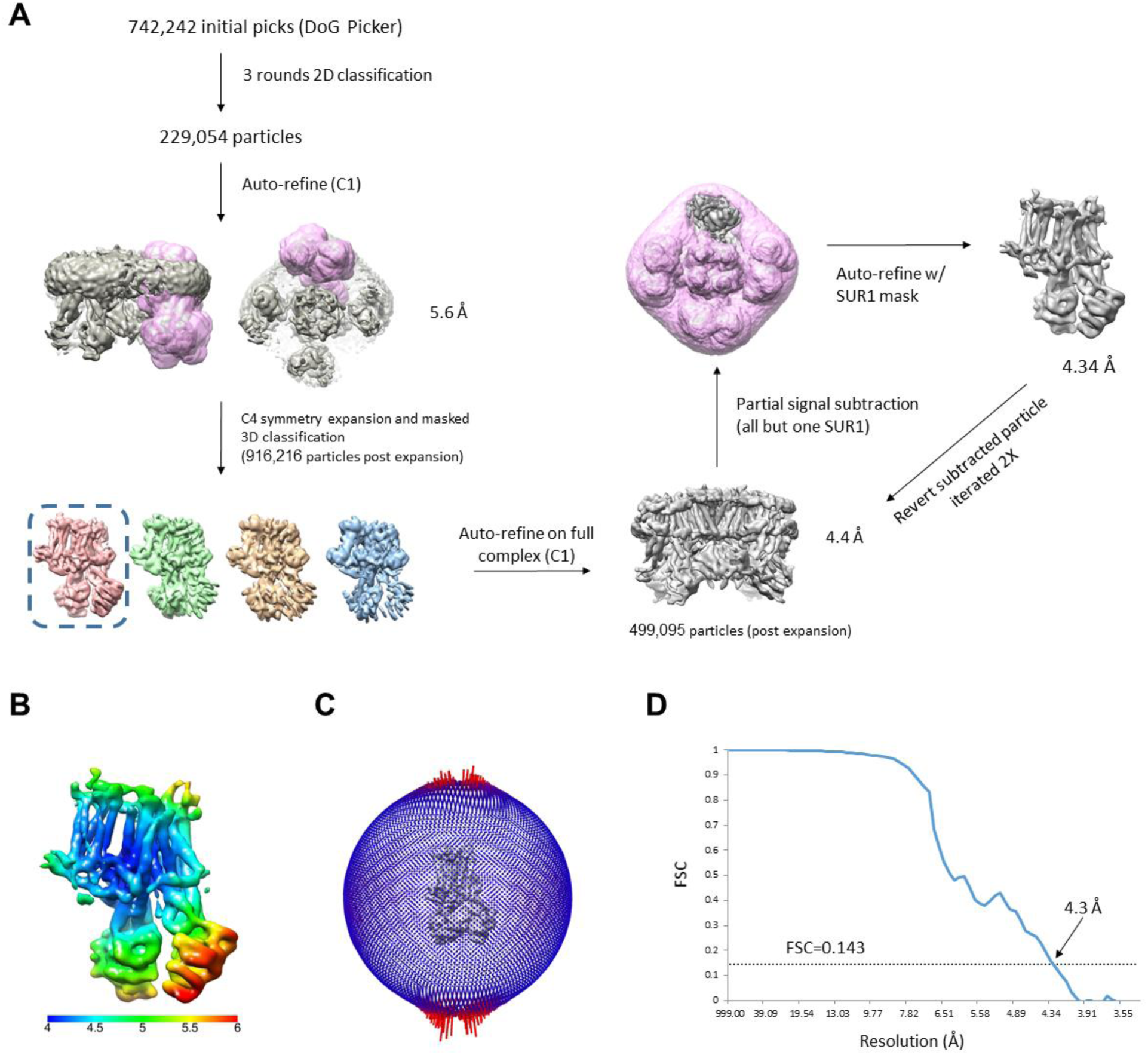
Data processing workflow for the CBZ/ATP state. **(A)** Overview of data processing workflow. Particle picking was performed automatically with DoGPicker and manual inspection. All other image processing steps were performed in RELION-3. **(B)** Local resolution plot of locally refined SUR1 map. **(C)** Angular distribution plot. **(D)** Fourier shell correlation (FSC) of two independent half maps of locally refined SUR1.

**Figure 1-figure supplement 3:**
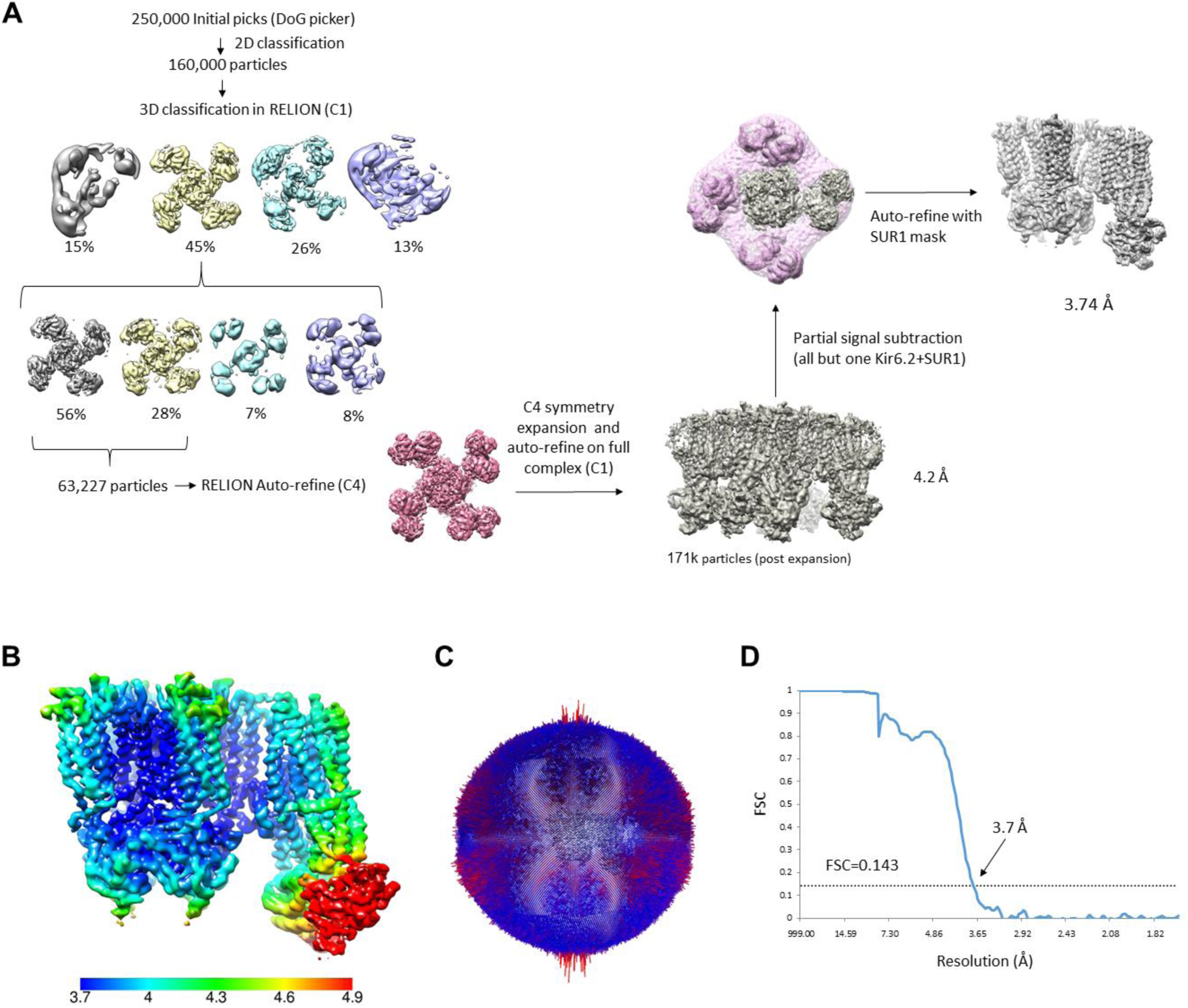
Data processing workflow for the GBC/ATP state. **(A)** Overview of data processing workflow. Note the dataset used was previously published in Martin et al. (34) and the particles that were included in the final reconstruction (EMD-7073) were used for focused refinement in RELION-3. **(B)** Local resolution plot of the locally refined SUR1 map. **(C)** Angular distribution plot. **(D)** Fourier shell correlation (FSC) of two independent half maps of locally refined SUR1.

**Figure 1-figure supplement 4:**
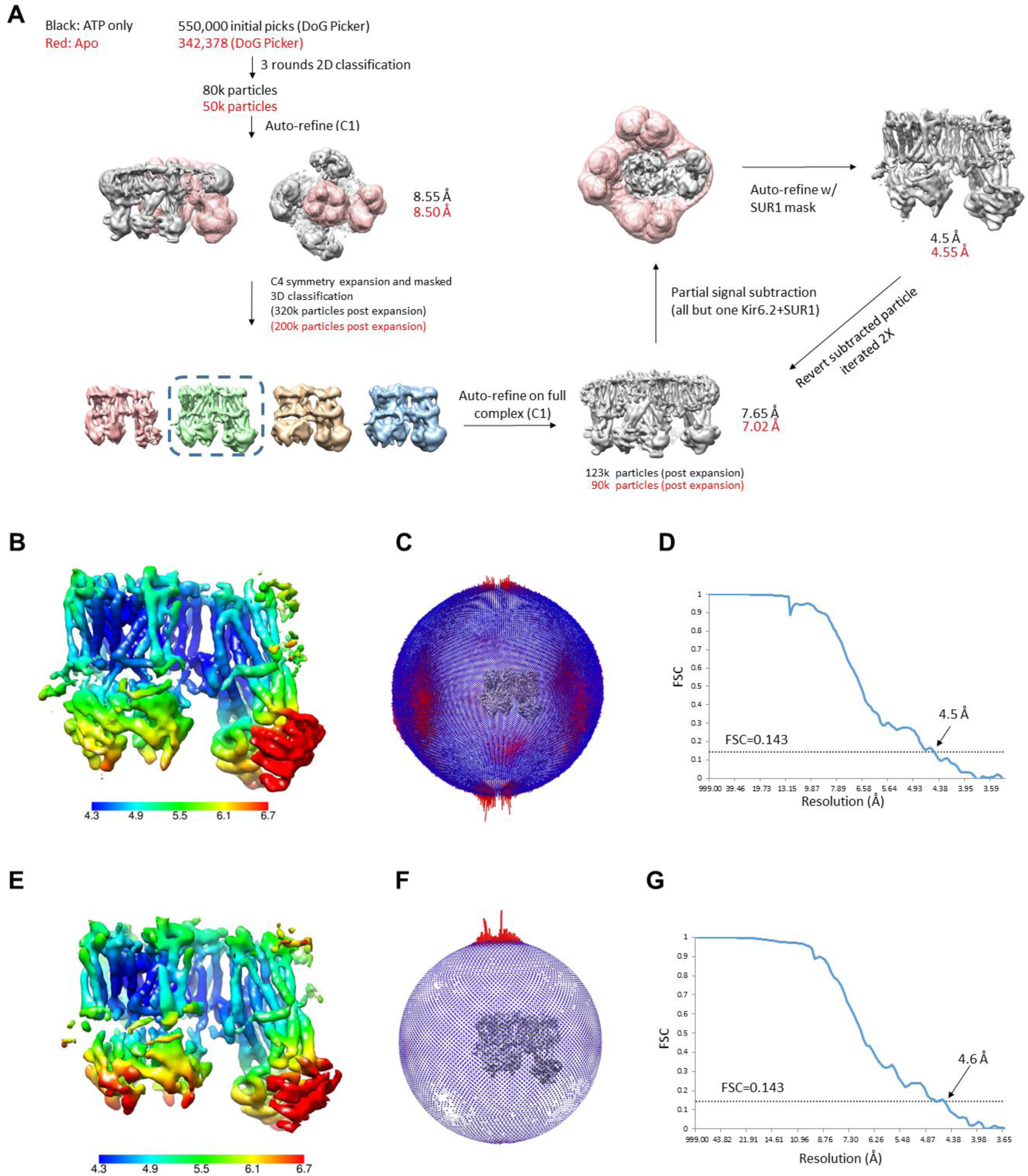
Data processing workflow for the ATP only state and the apo state. **(A)** Data processing flow for the ATP only state dataset. The Apo state dataset was processed in the same manner and relevant numbers are shown in red. **(B-D)** Local resolution map, angular distribution plot, and FSC plot for the ATP only dataset. **(E-G)** Local resolution map, angular distribution plot, and FSC plot for the apo state dataset.

**Figure 1-figure supplement 5:**
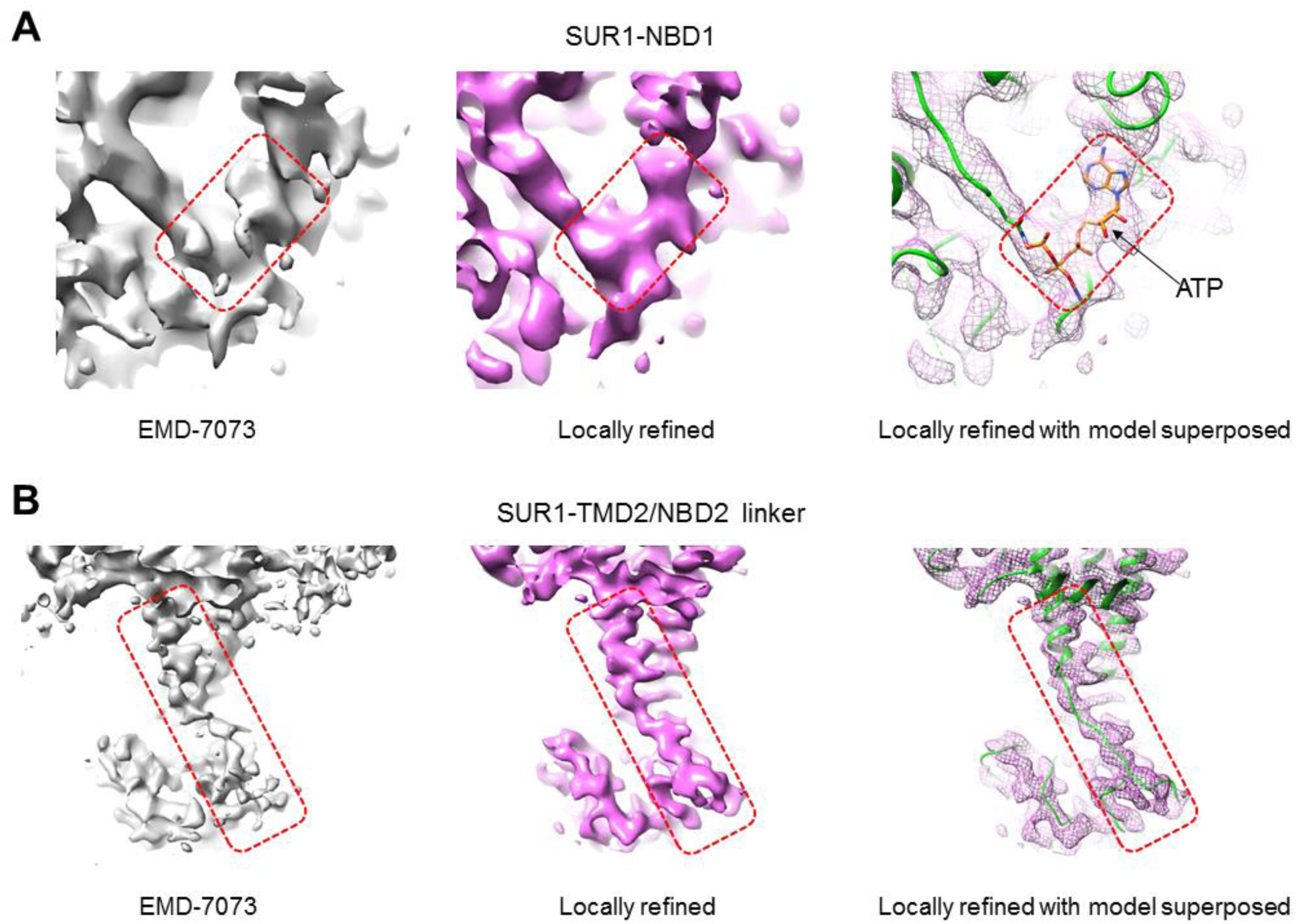
Comparison of the GBC/ATP state maps before and after focused refinement. **(A)** *Left*: Cryo-EM C4 map of the SUR1-NBD1 region published in Martin et al. (EMD-7073) contoured to 2.6σ; *middle*: the same region from the locally refined map contoured to 8.9σ; *right*: same as the middle panel except the density is displayed in mesh and the structural model of the protein and bound ATP are superposed. **(B)** Same as **(A)** but showing the linker region connecting TMD2 and NBD2.

**Figure 2-figure supplement 1:**
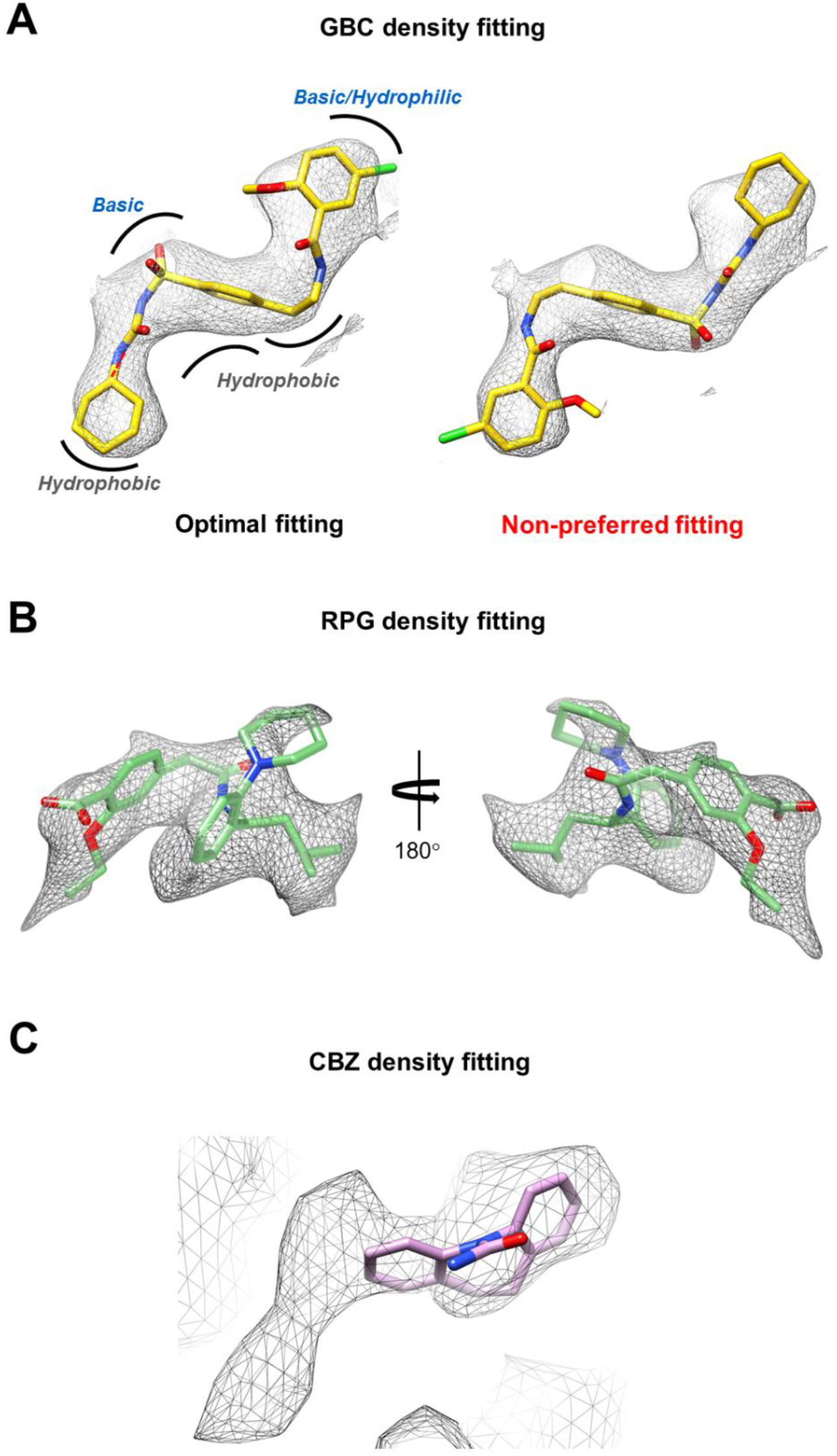
Density fitting for GBC, RPG, and CBZ. **(a)** CryoEM density of GBC from the focus-refined SUR1 map contoured to 8.5σ. *Left*: optimal fitting of the GBC structure into the density. The electrostatic nature of the residues in the binding pocket surrounding GBC (see Fig.3b) are shown to demonstrate general electrostatic mismatch with GBC if the molecule were modeled into the density in the flipped orientation by 180° shown on the r*ight*. **(b)** CryoEM density of RPG from the focus-refined SUR1 map contoured to 9σ with RPG fitted into the density. **(c)** CryoEM density of CBZ from the locally refined SUR1 map contoured to 8.5σ. A CBZ molecule is placed into one end of the density to illustrate that one stationary CBZ molecule cannot account for the full ligand density observed.

**Figure 3-figure supplement 1:**
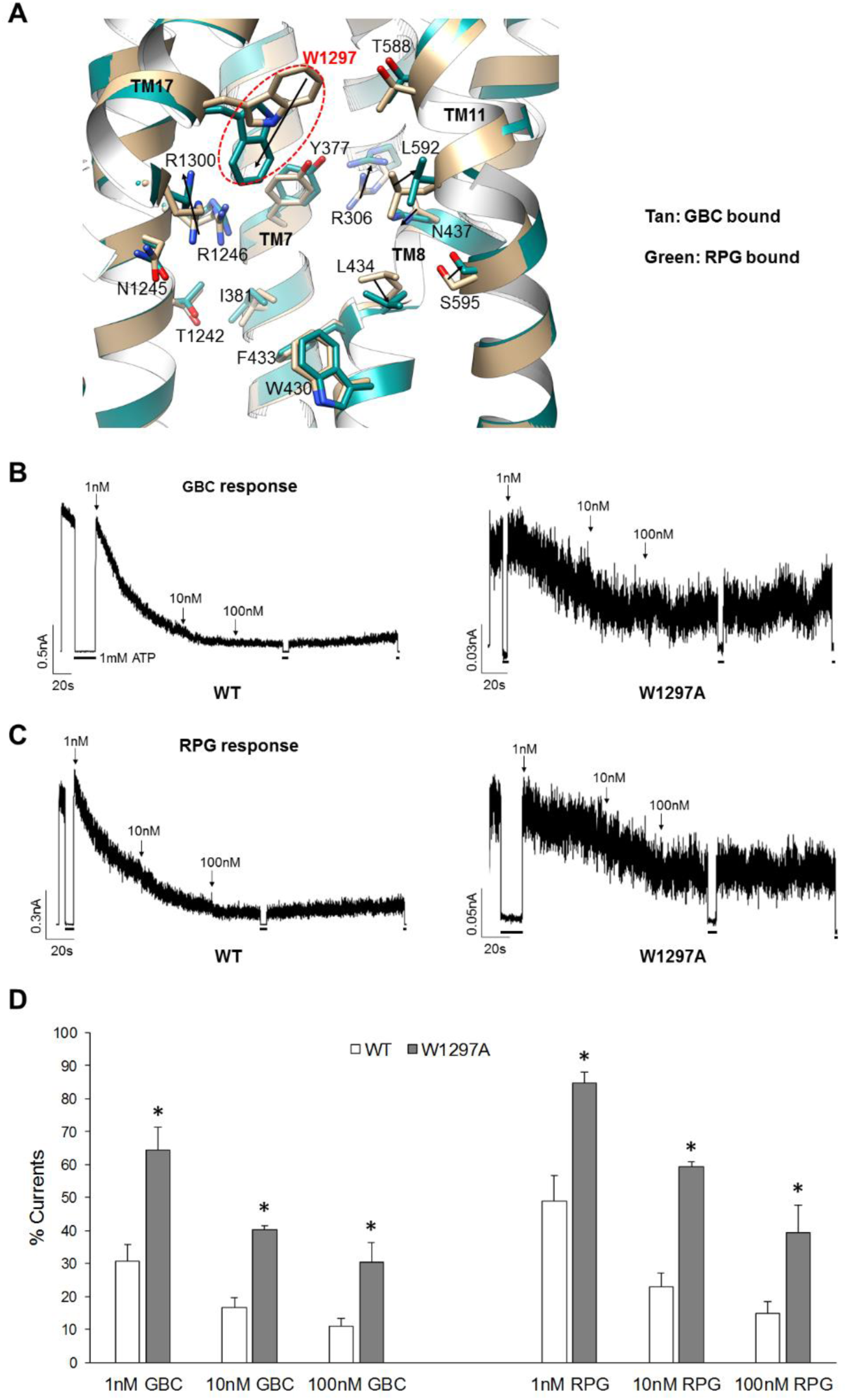
Comparison of SUR1 drug binding pocket structures in the GBC and RPG bound states. **(A)** Structural models showing the differences observed in the residues around GBC (in tan) or RPG (in green). Arrows indicate the direction of movement. W1297, which has its side-chain pointed in opposite directions in the two structures and is mutated to A for functional testing in **(B, C)** is circled and labeled in red. **(B, C)** Representative inside-out patch-clamp recording traces showing response of WT and the SUR1-W1297A mutant channels to 1, 10, 100nM GBC **(B)** or RPG **(C)**. Application of the drugs at the various concentrations are indicated by the arrows. Black bars on the bottom of the traces indicate exposure to 1mM ATP for baselines. **(D)** Histograms showing the effects of the W1297A mutation on channel sensitivity to GBC or RPG. Each bar represents mean s.e.m. of 4-5 patches. The asterisk indicates statistical significance compared to WT (*p* < 0.05) using unpaired, two-tailed Student’s *t*-test.

**Figure 4-figure supplement 1.**
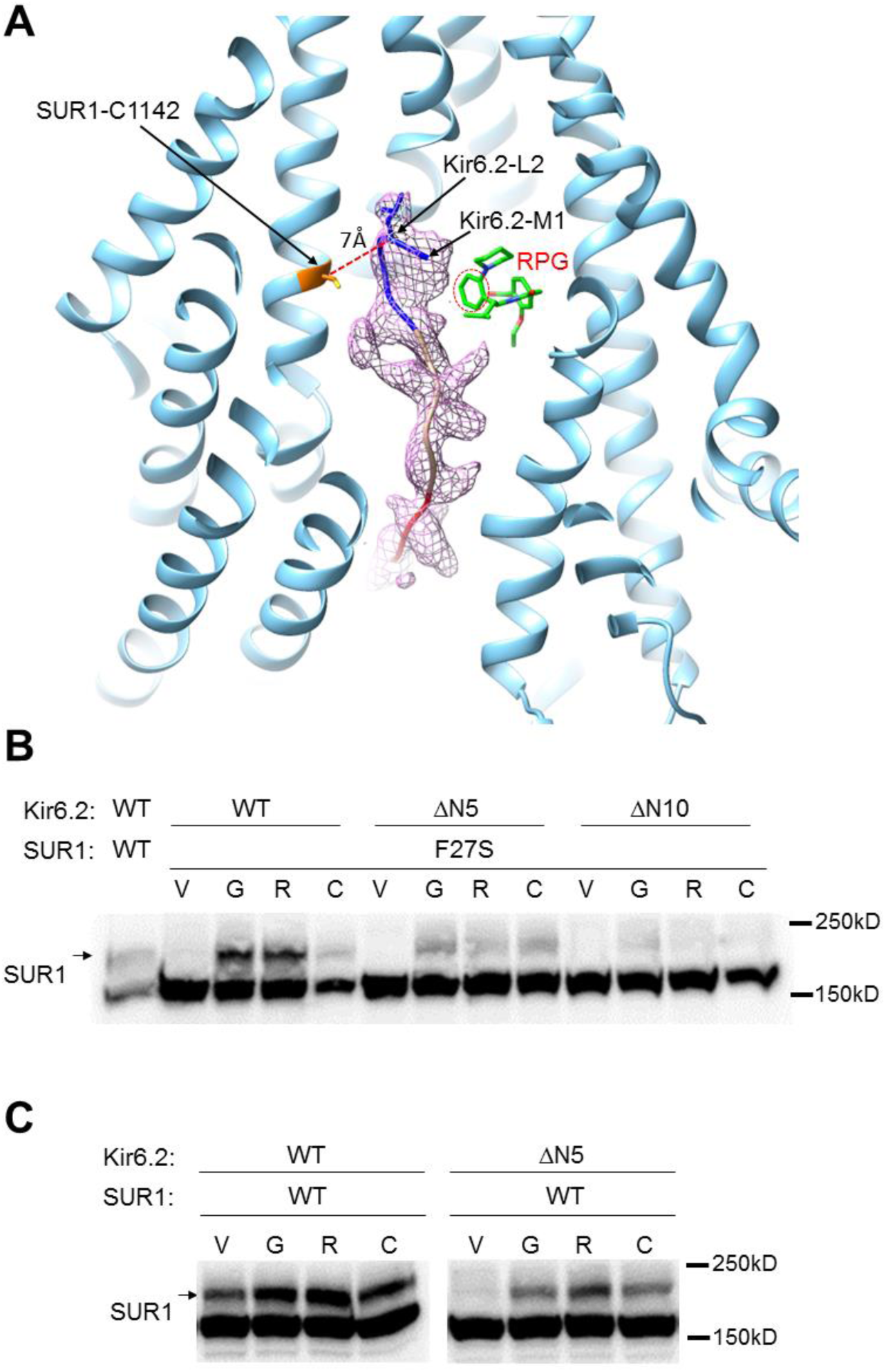
The distal N-terminus of Kir6.2 interacts with SUR1 and is required for channel biogenesis and pharmacochaperone rescue. **(A)** Kir6.2 N-term cryoEM density (pink mesh) with superposed polyalanine model shown in the RPG (green) bound SUR1 structural model. The piperidino moiety of RPG is highlighted with dotted red line to show its close proximity to the N-terminal methionine of the modeled Kir6.2 N-terminal peptide. The relationship between SUR1 C1142 and Kir6.2 L2 is shown to illustrate their close proximity, with Ca-Ca distance of ∼7Å. The Kir6.2 N-term density map was obtained by removing densities corresponding to modeled SUR1 and RPG from the focus-refined RPG/ATP map using the Color Zone option in Chimera, contoured to 12σ. The polyalanine model of the Kir6.2 N-term corresponding to amino acids 1-5 is shown in blue, 5-10 in tan, and 10-15 in red. **(B)** Western blot showing that deletion of Kir6.2 amino acids 2-5 (ΔN5) or amino acids 2-10 (ΔN10) attenuated or nearly abolished, respectively, the pharmacochaperoning effect [compared to 0.1% DMSO vehicle control (V)] of GBC (G; 1µM), RPG (R; 1µM), and CBZ (C; 10µM) on the SUR1-F27S mutant. **(C)** Western blot showing deletion of amino acids 2-5 of Kir6.2 also greatly impaired maturation of WT SUR1 (V), although GBC, RPG, and CBZ still have some albeit reduced effects on boosting maturation of SUR1.

**Figure 4-figure supplement 2.**
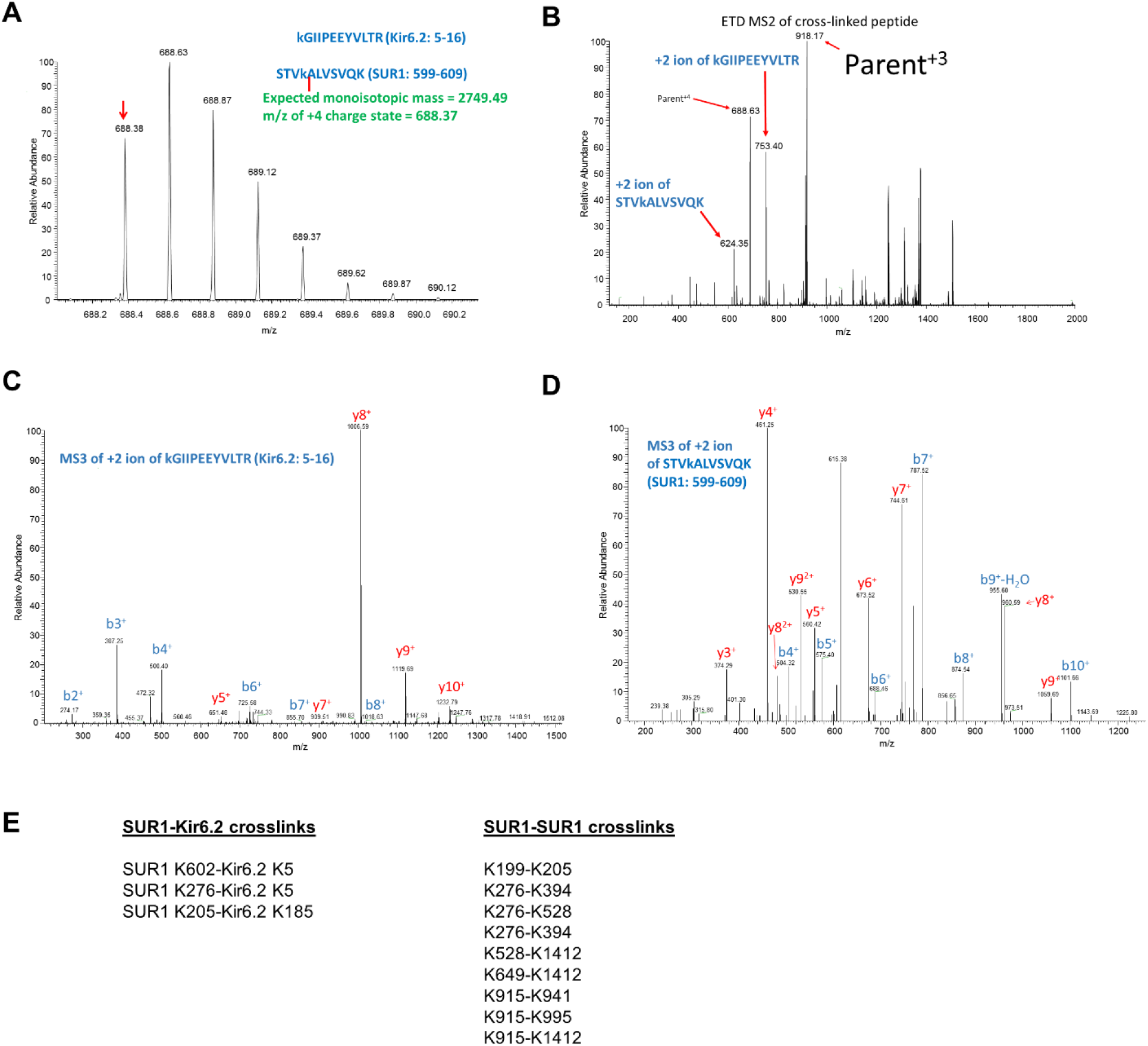
Mass spectrometric identification of a chemical crosslink between Kir6.2 peptide KGIIPEEYVLTR (5-16) and SUR1 peptide STVKALVSVQK (599-609). **(A)** MS survey scan measuring the mass of the +4 charge state of the crosslinked peptide with matching calculated and measured m/z values for the monoisotopic ion (m/z = 670.34, red arrow). **(B)** Electron transfer dissociation of crosslinked m/z =670.34 ion with liberated peptides indicated by red arrows following cleavage of the disulfide bond in the DSP crosslinker. The other major ions are the +4 parent ion and +3/+2 charged reduced parent ions that remained unfragmented. **(C and D)** MS^3^ spectra using collision-induced dissociation to confirm the identities of liberated peptides KGIIPEEYVLTR and STVKALVSVQK (red arrows in panel b). Matched b (blue) and y (red) fragment ions are indicated. **(E)** Crosslinked peptides identified from purified channels in the presence of 1µM glibenclamide.

## Abbreviations

CBZ: carbamazepine
CFTR: cystic fibrosis transmembrane conductance regulator
GBC: glibenclamide
K_ATP_: ATP-sensitive potassium channel
Kir6.2: inwardly rectifying potassium channel 6.2
NBD: nucleotide binding domain
PIP2: phosphatidylinositol 4,5-bisphosphate
RPG: repaglinide
SU: sulfonylureas
SUR1: sulfonylurea receptor 1
TM: transmembrane; TMD: transmembrane domain.

